# Human iPSC-derived renal cells change their immunogenic properties during maturation: Implications for regenerative therapies

**DOI:** 10.1101/2021.03.01.432225

**Authors:** Bella Rossbach, Krithika Hariharan, Nancy Mah, Su-Jun Oh, Hans-Dieter Volk, Petra Reinke, Andreas Kurtz

**Author notes:** Equal contribution (Petra Reinke, Andreas Kurtz).

## Abstract

Therapeutic success of human induced pluripotent stem cell (hiPSC)-based therapies critically depends on immunological compatibility of the hiPSC-derived transplant. As grafted hiPSC-derived cells are often immature, we hypothesized that their immunologic properties may change due to post-grafting maturation. Subsequently, this will affect their interaction with the host immune system and may compromise graft tolerance. In the present study allogeneic and autologous cellular immunity of primary cells, therof reprogrammed hiPSC, hiPSC-derived progenitor and terminally differentiated cells was investigated *in vitro* by using renal cells as a model system. In contrast to allogeneic primary cells, hiPSC-derived early renal progenitors and mature renal epithelial cells were both tolerated not only by autologous but also by allogeneic T cells. These immune-privileged properties resulted from active immune-modulation and low immune visibility, which declined during the process of cell maturation. However, autologous and allogeneic natural killer (NK) cell responses were not suppressed by hiPSC-derived renal cells and efficiently changed NK cell activation status. These findings clearly show a dynamic stage-specific dependency of autologous and allogeneic T- and NK cell responses to the hiPSC-derived renal cell lineage with consequences for effective cell therapies. The study suggests that hiPSC-derived early progenitors may provide advantageous immune suppressive properties when applied in cell therapy. The data furthermore indicate a need to suppress NK cell activation in allogeneic as well as autologous settings.

## 1. Introduction

Human induced pluripotent stem cells (hiPSC) provide an unlimited source material for functionally differentiated cells suitable in cell-replacement therapies (CRT) and tissue engineering.^[1,2]^ Compared to human embryonic stem cells (hESC), iPSC-technology offers the possibility of personalized autologous CRT and thus might overcome rejection barriers connected to alloimmunogenicity. Surprisingly, transplanted syngeneic murine iPSC (miPSC) were shown to be rejected in a T cell driven manner.^[3]^ Consequently, the immunological effects of clinically applicable hiPSC-derivatives are a major concern. Reprogramming and cultivation based neoantigens may cause some of these immunological effects, however, despite the development of integration-free reprogramming techniques and xeno-free media, iPSC-derived cells may still invoke a variable response from the immune system. ^[3,4,5]^ For example, certain miPSC-derivatives like endothelial cells, dermal cells, bone marrow cells, hepatocytes and neuronal cells were not rejected in syngeneic recipients, whereas transplanted miPSC-derived cardiomyocytes elicited significant levels of T cell infiltration.^[6–8]^ In a humanized mouse model, hiPSC-derived retinal pigment epithelial cells (RPEC) were tolerized by autologous reconstituted T cells, whereas differentiated smooth muscle cells (SMC) were rejected.^[9]^ It was demonstrated that the aberrant expression of Zymogen granule protein 16 (ZG16) was inducing the immunogenic nature of the hiPSC-derived SMC. Aberrant expression of Zg16, as well as of HORMA domain-containing protein 1 (Hormad1) has already been shown in autologous transplanted miPSC and deemed responsible for their immunogenic nature.^[3]^

For efficient CRT, cells at different maturation stages may be needed depending on patient, disease, tissue, desired cell type and the anticipated mode of action. Terminally differentiated cells such as hiPSC-derived retinal pigment epithelial cells (RPEC) might directly replace damaged cells in macular degenerative diseases, whereas further cell proliferation and differentiation post-grafting may be required to rebuild tissue structure, like for the restoration of the hematopoietic system.^[10,11]^ However, it remains unknown whether the maturation stage of hiPSC-derived cells modulates their immunogenic characteristics and immunomodulatory properties after transplantation, and subsequently inform clinicians about clinical suitability of developmental maturity and the associated need for immunological control.

In addition, due to the logistic and cost related challenges of autologous hiPSC-based therapies, the allogeneic off-the-shelf approach is a major focus of research. Efforts are underway to establish an unlimited resource of hiPSC-lines, which are haploidentical and homozygous for common human leucocyte antigen (HLA) alleles to decrease allogeneic mismatches of the differentiated products.^[12,13]^

We used the kidney as a model system to investigate maturation stage dependent auto- and allogenicity of hiPSC-derived cells. Renal cells are a relevant example for epithelial cells and transplantation of kidneys is a major clinical need. Different type of renal cells at defined developmental stages can be derived nowadays from hiPSC.^[14,15]^ These cells could therapeutically be used for CRT in patients suffering from acute kidney injury (AKI) or chronic kidney disease (CKD), global health problems with increasing prevalence.^[16]^ Indeed, transplanted miPSC and miPSC-derived renal progenitors, respectively, were shown to support regeneration processes and improve renal function in immunocompromised and immune-suppressed models of AKI, respectively.^[17,18]^

Induction of intermediate mesoderm cells (IMC) is the first step of renal differentiation from hiPSC, followed by epithelialization and specification of the nephron elements, including proximal epithelial cells (PTC). We analyzed the immunological responses of human T and natural killer (NK) cells towards hiPSC and hiPSC-derived IMC and PTC using sensitive *in vitro* assays. Damage of PTC is the leading cause of AKI and subsequently CKD. Moreover, we collected primary urinary cells (pUC) from the hiPSC-donors to directly compare the donor-specific allogeneic and autologous response of hiPSC-derived and non-hiPSC-derived cells in an isogenic setup. Immune-phenotypic analysis revealed decreased HLA-ABC and HLA-DR expression in hiPSC and hiPSC-derived renal cells compared to pUC. Although allogeneic T cell activation was observed against pUC, neither autologous nor allogeneic hiPSC or hiPSC-derived renal cells induced T cell responses. However, hiPSC and hiPSC-derived renal cells showed susceptibility to NK cells. Active immunomodulatory properties were observed in hiPSC, IMC and early stage PTC, which may explain the attenuated immune response of the cells even in allogeneic conditions and implicate an at least temporary immune-privileged status of hiPSC and hiPSC-derived precursor cells. Taken together, we identified an immune-privileged status of hiPSC-derived IMC and PTC, which declines with cell maturation.

## Results

### 2.1. Generation and maintenance of hiPSC-derived renal cells

Isolated primary cells from urinary sediments of healthy donors include mostly exfoliated renal tubular cells and urinary tract epithelial cells.^[19]^ Expanded pUC showed heterogeneous morphologies consisting of cuboid and spindle-like cells and were used to generate the urinary-cell derived hiPSC-lines BCRTi004-A and BCRTi005-A (**Figure 1**a).^[20,21]^ Differentiation of BCRTi004-A and BCRTi005-A consistently generated IMC and PTC using a step-wise protocol (Figure 1a, b).^[14]^ Typical cobblestone morphologies were observed in PTC and maintained in long-term cultivated proximal tubular cells (LT-PTC) at least for 26 days (Figure 1b). Analysis of cell-type specific marker expression by flow cytometry revealed up to 80 % differentiation efficiencies (Figure 1c). IMC were defined by expression of the IM-specific markers LIM homeobox 1 (LHX1) and Paired box 2 (PAX2). PTC were assessed by occurrence of Aquaporin 1 (AQP1) and Sodium-potassium-adenosine triphosphatase (Na/K-ATPase), which remained stably expressed after continued cultivation for 14 days on Geltrex (Figure 1b, c and **Figure S1**). Additional immunostaining revealed co-localization of LHX1 and PAX2 in IMC, as well of AQP1 and Na/K-ATPase, respectively, in PTC and LT-PTC (**Figure S2**). Principal component analysis (PCA) of RNA-sequencing data of hiPSC, hiPSC-derived renal cells and pUC revealed differential clustering of the respective cell types (Figure 1d).

**Figure 1:**
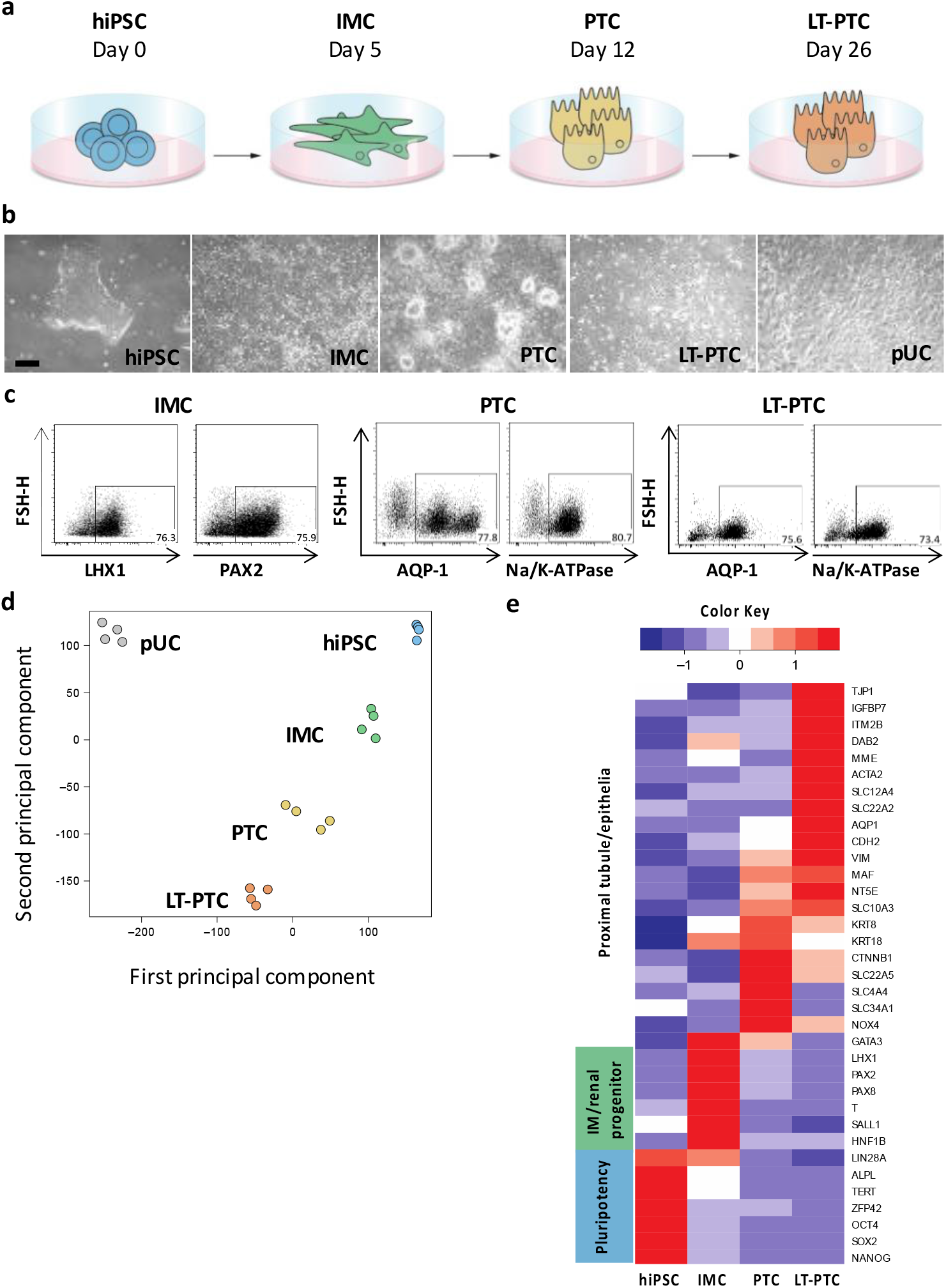
Reprogrammed pUC differentiate with high efficiency into renal progenitors and into PTC, which can be stably maintained *in vitro*. **a)** hiPSC, generated from pUC, were differentiated into IMC and PTC using a step-wise protocol. Differentiated PTC were further cultivated for two additional weeks (LT-PTC). **b)** Differentiated cells were examined by phase-contrast microscopy and showed stage specific cell morphology. **c)** Cell stage specific proteins for IMC, PAX2 and LHX1, and for PTC, AQP1 and Na/K-ATPase, were analyzed by flow cytometry to determine differentiation efficiencies. Transcriptome analysis revealed **d)** differential clustering as depicted in a PCA plot and **e)** stage specific gene expression in the primordial pUC, hiPSC and hiPSC-derived IMC, PTC and LT-PTC. Data are representative of biological duplicates per cell type and per cell line. The expression values (FPKM) of each gene (row) are normalized by a row z-score. Scale bar is equivalent to 100 μm.

For more in-depth analysis of differentiation and maturation progression, stage-specific marker gene expression was compared. The data confirmed decreased expression of pluripotency-associated genes like Octamer-binding transcription factor 4 (OCT4), Sex determining region Y-box 2 (SOX2) and Telomerase reverse transcriptase (TERT) in IMC, PTC and LT-PTC. The markers LHX1, PAX2, PAX8, Gata binding protein 3 (GATA3) as well as the mesendoderm marker Brachyury (T) were specifically upregulated in IMC. Increased expression of PTC markers like the membrane transport proteins Solute carrier family 10 member 3 (SLC10A3), SLC12A4, AQP1, the epithelial markers Catenin beta 1 (CTNNB1), Keratin 8 (KRT8), KRT18 and the adhesion molecule Cadherin 2 (CDH2) occurred in the differentiated PTC and showed stable expression during long-term cultivation in LT-PTC (Figure 1e). Together, these results indicate that the hiPSC-derived renal cells faithfully recapitulate the stages IMC and PTC during kidney development and further maturation progression in the LT-PTC. In comparison, undifferentiated hiPSC did not show expression of IMC- and PTC-specific markers (Figure S3).

### 2.2. Immune-phenotype of hiPSC and hiPSC-derived renal cells

Expression of MHC class I (HLA-ABC) and MHC class II (e.g. HLA-DR) is essential for the recognition of antigens by cluster of differentiation (CD)8^+^ T cells and CD4^+^ T cells, respectively. The presence of HLA-ABC and HLA-DR molecules on the surface of hiPSC, IMC, PTC, LT-PTC and pUC was assessed to elucidate their capacity to elicit T cell responses. The cells were stimulated by interferon gamma (IFNγ), which induces upregulation of HLA-ABC and HLA-DR expression in pro-inflammatory environments triggered by infiltrating leukocytes, to simulate damaged tissue usually faced by cells in CRT.^[22]^ Under homeostatic conditions hiPSC, IMC and PTC expressed very low levels of HLA-ABC (**Figure 2**a). After stimulation with IFNγ, HLA-ABC expression was strongly upregulated in the hiPSC, IMC and PTC. HLA-DR was not detectable on these cell types even after IFNγ treatment (Figure 2b). Interestingly, LT-PTC in comparison showed increased HLA-ABC expression already under homeostatic conditions and expression levels further increased upon IFNγ stimulation. Additionally, HLA-DR expression in LT-PTC was induced by IFNγ stimulation. As expected, pUC expressed high levels of HLA-ABC, which was further elevated upon IFNγ stimulation and HLA-DR expression was induced too.

**Figure 2:**
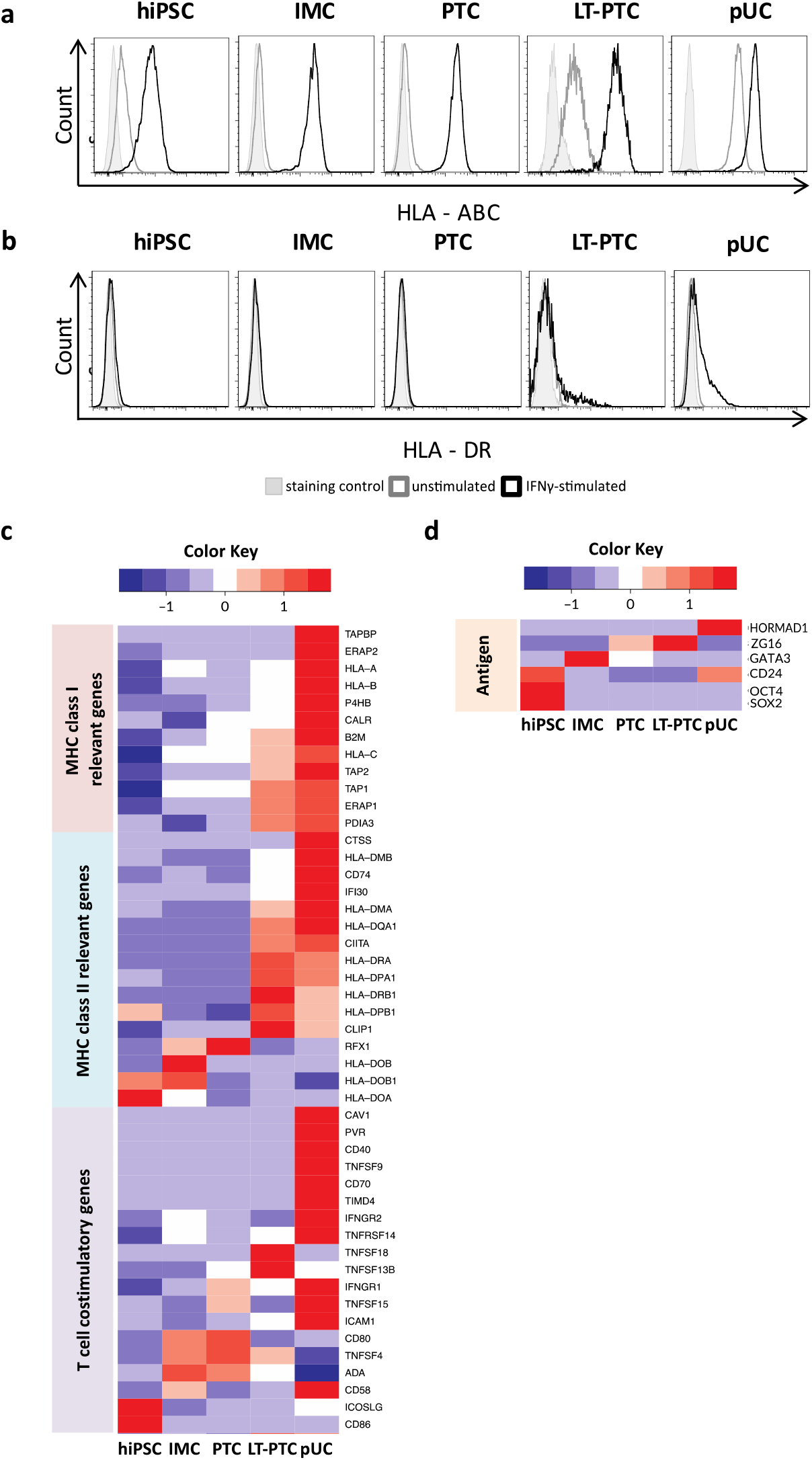
Immune-phenotype of hiPSC and hiPSC-derived renal cells are less mature compared to pUC, but is regulated by a pro-inflammatory environment. hiPSC, differentiated IMC, PTC and LT-PTC were analyzed for HLA-ABC and HLA-DR molecules on the cell surface by flow cytometry. Adult somatic pUC were used in comparison. Representative dot plots are shown for **a)** HLA-ABC and **b)** HLA-DR under homeostatic and under pro-inflammatory conditions induced by IFNγ stimulation. HLA-ABC molecules were low on the surface of hiPSC, IMC and PTC but were increased in LT-PTC, whereas pUC expressed moderate levels. HLA-DR molecules were only detectable in LT-PTC and pUC after IFNγ stimulation. **c,d)** Transcriptome profiles were used to analyze expression of **c)** MHC class I, MHC class II, T cell co-stimulatory molecules and **d)** common immunogenic antigens in hiPSC, and hiPSC-derived renal cells and expression intensities were compared to somatic pUC. Data are representative of three IFNγ stimulated biological replicates per cell type. The expression values (FPKM) of each gene (row) are normalized by a row z-score.

To gain a better understanding of the immune-phenotype of IFNγ-treated hiPSC, IMC, PTC, LT-PTC and pUC, transcriptomes were analyzed for the expression of genes related to MHC class I, MHC class II and T cell co-stimulatory factors (Figure 2c). MHC class I related genes like the respective polymorphic α-chains, the non-polymorphic β2-microglobulin (β2M), the transporter associated with antigen processing (TAP) TAP1 and TAP2 were expressed at lower levels in hiPSC compared to pUC and upregulated with progressing differentiation. Immune-maturation continued in LT-PTC after specification of the proximal tubular phenotype in PTC. MHC class II genes showed variable expression throughout the developmental stages. However, Class II major histocompatibility complex transactivator (CIITA), crucial for MHC class II expression, was detectable in LT-PTC and pUC only. Common T cell co-stimulatory molecules like CD80 and CD86, were detectable at low levels in hiPSC, IMC, PTC, LT-PTC as well as in pUC. Other co-stimulatory factors like CD40, CD70, Tumor necrosis factor ligand superfamily member 9 (TNFSF9) showed the highest expression in pUC, whereas the expression in hiPSC and renal derivatives was markedly low and did not show differentiation stage associations.

Next, we examined the transcript expression of potentially immunogenic antigens like SOX2, OCT4, HORMAD1, ZG16, CD24 and GATA3, which were previously described to elicit immune responses leading to the rejection of miPSC, hiPSC and their derivatives by antigen-specific T cells in preclinical models (Figure 2d).^[4,23–25]^ OCT4 was highly expressed in hiPSC, while residual OCT4 expression was strongly reduced in IMC, PTC and LT-PTC. CD24 was highly expressed in hiPSC, showing downregulation during the course of renal differentiation and expression almost disappeared in LT-PTC. The transcription factor GATA3 is selectively expressed during embryogenesis of the human kidney and thus highly transcribed in IMC.^[26]^ HORMAD1 was not detectable at any stage during renal differentiation and was barely detectable in pUC, whereas ZG16 was marginally detected in PTC and LT-PTC.

In summary, donor-identical pUC are more immunogenic than hiPSC and hiPSC-derived renal cell types, however, phenotypic immunogenicity moderately increases at later cellular states in LT-PTC. Moreover, potentially immunogenic antigens show expected cell type specific expression patterns.

### 2.3. Autologous T cell response against hiPSC-derived renal cells

Although the analysis of the immune-phenotype of hiPSC-derived renal cells revealed low expression of genes of the MHC class I and MHC class II complexes compared to primary somatic cells, prediction of the immunogenicity based on transcript and protein expression pattern alone is not possible as immune response can be triggered by very low levels of HLA/peptide complexes. *In vitro* one-way mixed lymphocyte reactions (one-way MLR) were performed to elucidate T cell responses triggered by autologous hiPSC and hiPSC-derived renal cells. Peripheral blood mononuclear cells (PBMC) isolated from the pUC-donors, used for the generation of hiPSC-lines BCRTi004-A and BCRTi005-A, respectively, were used (**Figure 3**a). Thus, all stimulator cells (hiPSC, pUC) shared identical HLA-genotypes. CD4^+^ and CD8^+^ T cell proliferation was monitored after co-cultivation of fluorescently labeled autologous PBMCs with hiPSC, IMC, PTC, LT-PTC and pUC, respectively (Figure 3b). A negligible fraction of CD4^+^ and CD8^+^ T cells proliferated in response to stimulation by any of the autologous hiPSC-derived renal cell types or undifferentiated hiPSC (Figure 3c). In summary, renal cells differentiated from hiPSC did not show susceptibility to autologous T cells (Figure 3d).

**Figure 3:**
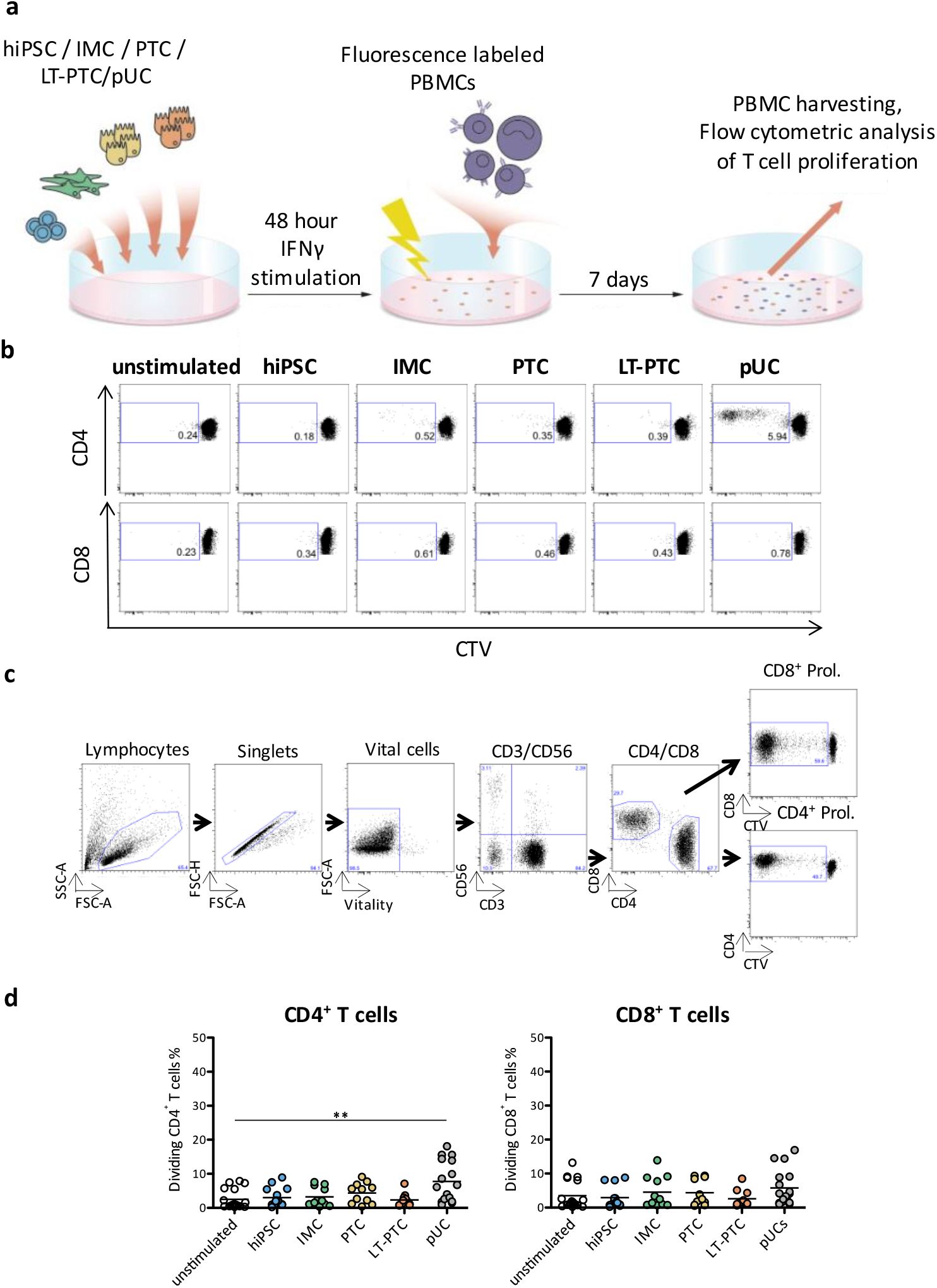
Autologous derived IMC, PTC and LT-PTC do not elicit T cell proliferation. Co-cultures of IFNγ stimulated hiPSC, IMC, PTC, LT-PTC and pUC with autologous PBMCs, respectively, were performed for 7 days and T cell proliferation was tracked using a fluorescence based method as shown in **a)**. Proliferation of CD4^+^ and CD8^+^ T cells was assessed using flow cytometry and gating was performed as depicted in **b)** as shown for SEB stimulated sample. **c)** Representative plots indicate absence of proliferation of CD4^+^ and CD8^+^ T cells, respectively, when stimulated with autologous hiPSC and hiPSC-derived IMC, PTC, LT-PTC and pUC. **d)** Graph depicts summary of 11-18 independent experiments showing the mean. Statistical analysis was performed using one-way ANOVA followed by Dunn’s post-test. *p < 0,05; **p < 0,01; ***p < 0,001.

### 2.4. Allogeneic T cell response against hiPSC-derived renal cells

Allogeneic off-the-shelf hiPSC-lines could represent an attractive source for clinical application. To assess the immunogenicity of allogeneic hiPSC and hiPSC-derived renal cells, PBMCs donated by unrelated unmatched healthy donors were co-cultured with hiPSC, IMC, PTC, LT-PTC and pUC, respectively, in one-way MLRs (Figure 3a, b). Available HLA-types of allogeneic PBMC from healthy donors showed at most one shared HLA-A/B/DR-allele with the hiPSC-lines (Figure S4). Tracking of CD4^+^ and CD8^+^ T cells in PBMC showed high proliferation response against allogeneic pUC (**Figure 4**a, b). Analysis of involved T cell subpopulations revealed that pre-formed memory T cells as well as naïve T cells reacted against allogeneic pUC (Figure 4c, Figure S5). In contrast, although HLA-identical with the allogeneic pUC, hiPSC, IMC, PTC and LT-PTC did not induce allogeneic T cell proliferation in any of the PBMC originating from unmatched healthy individuals. Further expression analysis of activation marker HLA-DR on CD4^+^ and CD8^+^ T cells after 7 days of co-cultivation confirmed the reduced immunogenicity of allogeneic hiPSC and renal descendants compared to allogeneic pUC (Figure S6). Furthermore, the level of released pro-inflammatory tumor necrosis factor alpha (TNFα) on day 3 was only elevated after co-cultivation with allogeneic pUC (Figure S7). In clinical situations, patients undergoing kidney CRT may present increased memory T cell levels and variability. Diabetic patients have increased numbers of allogeneic memory T cells compared to healthy individuals.^[27]^ We therefore used PBMCs from patients with diabetic nephropathy to study the rejection characteristics of allogeneic hiPSC-derived renal cells, and pUC (**Table S**1). Although pUC again induced strong T cell proliferation, hiPSC and hiPSC-derived renal cells sharing the HLA-type with the pUC did not (Figure 4d). Furthermore, direct comparison of all experimental groups confirmed that the T cell responses between autologous and allogeneic hiPSC, IMC, PTC and LT-PTC are essentially indistinguishable in comparison to pUC (Figure S8). In conclusion, hiPSC and hiPSC-derived renal cells showed immune-privileged properties. They neither induced proliferation and activation of allogeneic naïve T cells nor of pre-formed allogeneic memory T cells, while HLA-identical pUC elicited robust allogeneic T cell responses.

**Fig. 4:**
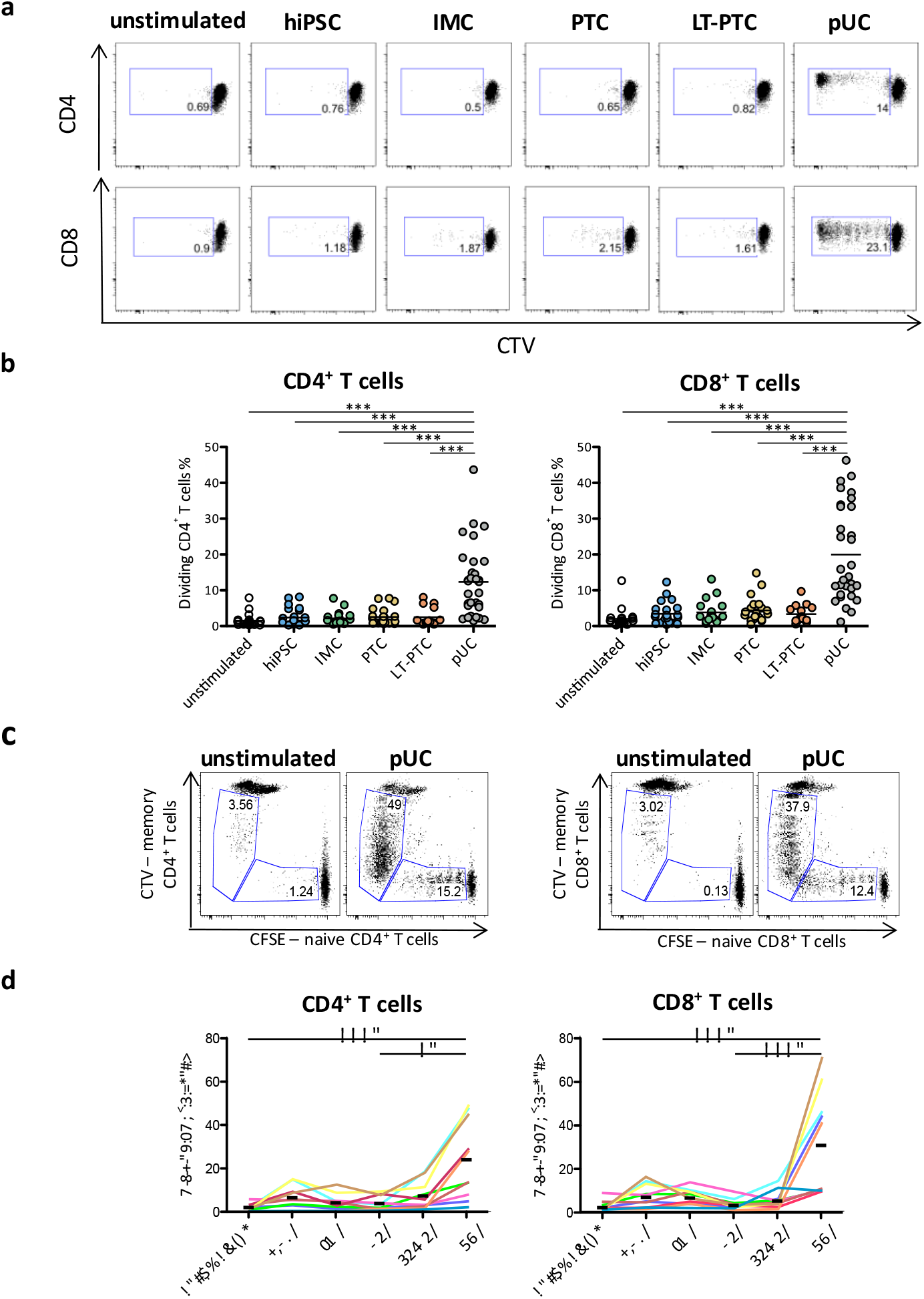
HLA-mismatched, allogeneic hiPSC, IMC, PTC and LT-PTC fail to induce proliferation of T cells from healthy donors and patients with diabetic nephropathy, despite the pre-existence of allogeneic memory T cells in the periphery. **a)** Healthy donors were analyzed for the pre-existence of allogeneic memory T cells against primordial pUC. Before co-culture of PBMCs with pUC, naïve T cells were separated and labeled with CTV, whereas residual PBMCs contained among others memory T cells were stained with CFSE. Independent tracking of naïve T cells and memory T cells after seven days of co-culture using flow cytometry revealed proliferation of naïve and memory T cells in parallel as depicted**. b)** Representative dot plots did not indicate induction of T cell proliferation by allogeneic hiPSC, IMC, PTC and LT-PTC, whereas allogeneic HLA-identical primordial pUC elicited strong CD4^+^ and CD8^+^ T cell proliferation. **c)** Summary of performed co-cultures of hiPSC, hiPSC-derived renal cells and pUC with PBMCs of healthy donors for 20 up to 31 independent experiments. **d)** PBMCs of patients with diabetic nephropathy were co-cultured with allogeneic hiPSC, IMC, PTC, LT-PTC and pUC. Significant T cell proliferation was obtained against pUC, whereas hiPSC and hiPSC-derived renal cells did not induce allogeneic T cell proliferation. Each colored line (n=10) represents data from a unique co-culture experiment of patients PBMCs with one set of hiPSC, renal differentiated hiPSC and corresponding pUC. **c,d)** Statistical analysis was performed using one-way ANOVA followed by Dunn’s post-test. *p < 0,05; **p < 0,01; ***p < 0,001

### 2.5. Immunomodulatory properties of hiPSC, IMC, PTC and LT-PTC

To analyze the nature of the disabled allogeneic T cell response, active immunomodulatory properties were studied in hiPSC and hiPSC-derived renal cell types. Thus, hiPSC, IMC, PTC, LT-PTC, and pUC were added, respectively, as third party to allogeneic MLR. Activated B cells expressing CD40, CD80 and CD86 were used as allogeneic stimulators (Figure S9). Activated B cells induced an average of 40% responder CD4^+^ and CD8^+^ T cell proliferation (**Figure 5**a). Addition of hiPSC highly suppressed allogeneic CD4^+^ and CD8^+^ T cell proliferation. Furthermore, addition of IMC and PTC, respectively, also reduced T cell proliferation after allogeneic B cell stimulation, but to a lower extent than hiPSC. In comparison, LT-PTC did not exhibit immunomodulatory function, while the addition of allogeneic pUC to the allo-MLR even increased allogeneic T cell proliferation.

**Figure 5:**
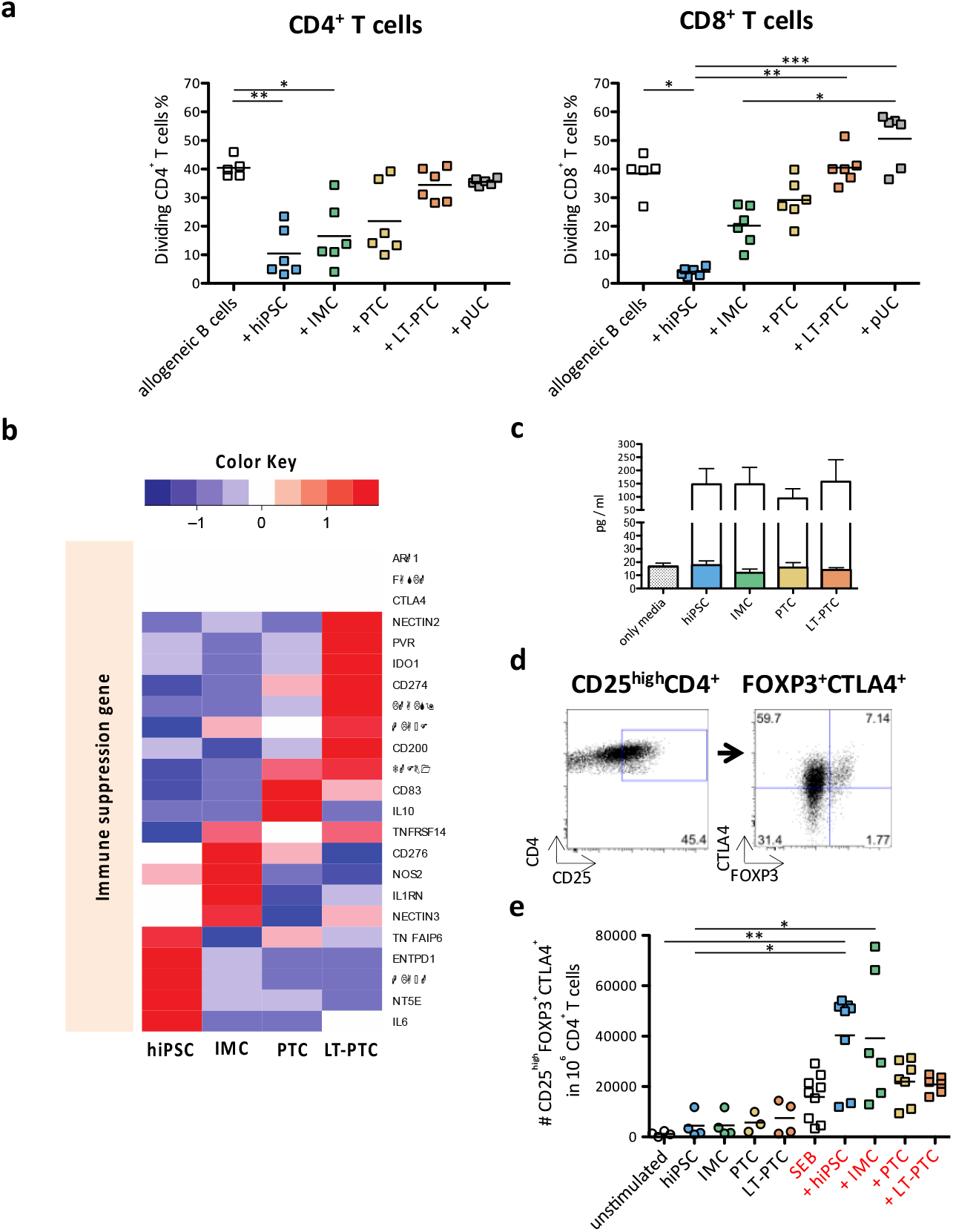
hiPSC, IMC and PTC but not LT-PTC exhibit T cell immunosuppressive properties. **a)** HLA-mismatched hiPSC, IMC, PTC, LT-PTC and pUC were added as third party to PBMCs stimulated with allogeneic B cells. Allogeneic T cell proliferation was inhibited by hiPSC, IMCs and PTCs. In comparison, LT-PTCs and pUC failed to suppress allogeneic B cell stimulated T cell proliferation. **b)** Transcriptome profiles of hiPSC and hiPSC-derived renal cells were analyzed for the expression of common immunosuppressive molecules and expression intensities were compared within the different cell types. Data are based on three IFNγ stimulated biological replicates for each cell type. **c)** ELISA were performed to quantify TGFβ proteins secreted by hiPSC and hiPSC-derived renal cells. Elevated levels of latent forms of TGFβ were detected in the supernatants of hiPSC, IMC, PTC and LT-PTC (colored bars indicate active TGFβ, while white bars indicate latent TGFβ protein). **d)** hiPSC and hiPSC-derived renal cells were analyzed for the capacities to polarize conventional CD4^+^ into a regulatory phenotype. After co-culture of PBMCs with hiPSC, IMC, PTC, LT-PTC and pUC, respectively, regulatory T cells were identified using flow cytometry as CD25^high^FOXP3^+^CTLA4^+^. **e)** Total number of CD4^+^ T cells with a regulatory phenotype was assessed after co-culture of PBMCs with hiPSC / hiPSC-derived renal cells alone, respectively, and after the addition of the polyclonal T cell stimulator SEB (n=4-9). For the statistical analysis one-way ANOVA was performed with subsequent Dunn’s post-testing. *p < 0,05; **p < 0,01; ***p < 0,001. The expression values (FPKM) of each gene (row) is normalized by a row z-score.

We used transcriptome data to identify potential candidates of immunosuppressive molecules secreted by hiPSC and hiPSC-derived renal cell (Figure 5b). Remarkably, we did not detect in hiPSC and hiPSC-derived renal cells at the mRNA level the common immunosuppressive molecules Arginase 1 (ARG1) and Fas ligand (FASLG), which were previously described to be expressed in PSC and inducing tolerance against allogeneic immunity.^[28,29]^ Also, expression of T cell inhibitory receptor cytotoxic T-lymphocyte associated protein 4 (CTLA4) and anti-inflammatory cytokine interleukin 10 (IL-10) was not observed. Instead, transcriptome data revealed association of renal cell maturation with RNA levels of the known immune suppression and immune escape genes Transforming growth factor beta 1 (TGFβ), Indolamin-2,3-dioxygenase (IDO1), CD274, the non-classical MHC class I molecules HLA-E as well as HLA-F and the ligands Poliovirus receptor (PVR) and Nectin cell adhesion molecule 2 (NECTIN2).

TGFβ is a pleiotropic polypeptide regulating multiple physiological processes including T cell growth and development.^[30]^ It has been demonstrated that TGFβ acts as a potent inducer of Forkhead box P3 (FOXP3), leading to the *de novo* generation of induced regulatory T cells (iTreg).^[31]^ Using enzyme-linked immunosorbent assay (ELISA), we confirmed protein secretion of TGFβ by hiPSC, IMC, PTC and LT-PTC. However, the obtained data also demonstrated that secreted TGFβ was still in its latent and thus inactive form (Figure 5c). We further wanted to investigate if the increased expression of latent TGFβ by hiPSC and hiPSC-derived renal cells in an allogeneic setting could lead to the conversion of conventional CD4^+^CD25-T cells into iTreg.

After the stimulation of PBMCs with allogeneic hiPSC, IMC, PTC and LT-PTC, respectively, the total number of CD4^+^ iTreg cells marked by the co-expression of CD25^high^CTLA4^+^FOXP3^+^ was identified (Figure 5d). We observed no increased numbers of FOXP3^+^ Tregs in any of the co-culture experiments (Figure 5e). In contrast, the additional presence of the polyclonal T cell activator Staphylococcal enterotoxin B (SEB) in one-way MLR with hiPSC and IMC, respectively, led to significantly higher numbers of CD25^high^CTLA4^+^FOXP3^+^ cells to SEB alone or unstimulated controls. In conclusion, hiPSC, IMC and PTC possess active immunomodulatory capacities. The immunosuppressive impact declines with progression of differentiation and LT-PTC and adult pUC did not exhibit anti-proliferative effects on stimulated allogeneic T cells.

### 2.6. Autologous and allogeneic NK cell responses to hiPSC and hiPSC-derived renal cells

Previous reports described contradictory results about the susceptibility of pluripotent stem cells to NK cells.^[4,32,33]^ We therefore examined the sensitivity of NK cells to hiPSC and the hiPSC-derived renal cells. NK cells express several stimulatory receptors, such as Natural killer group 2 (NKG2D) receptors, which promote cytotoxic and inflammatory activation and might lead to the elimination of ligand expressing cells.^[34]^ Transcriptome analysis of hiPSC, IMC, PTC, LT-PTC and pUC revealed expression of NKG2D ligands stress-related MHC class I polypeptide-related sequence (MIC) A, MICB, UL16 binding protein (ULBP) 2 and ULBP3, with highest expression in pUC (**Figure 6**a). Expression of NK cell inhibitory ligands like HLA-E, HLA-G, CD200 and C-type lectin domain family 2 member D (CLEC2D), however, were increased in hiPSC and in LT-PTC in comparison to pUC.

**Figure 6:**
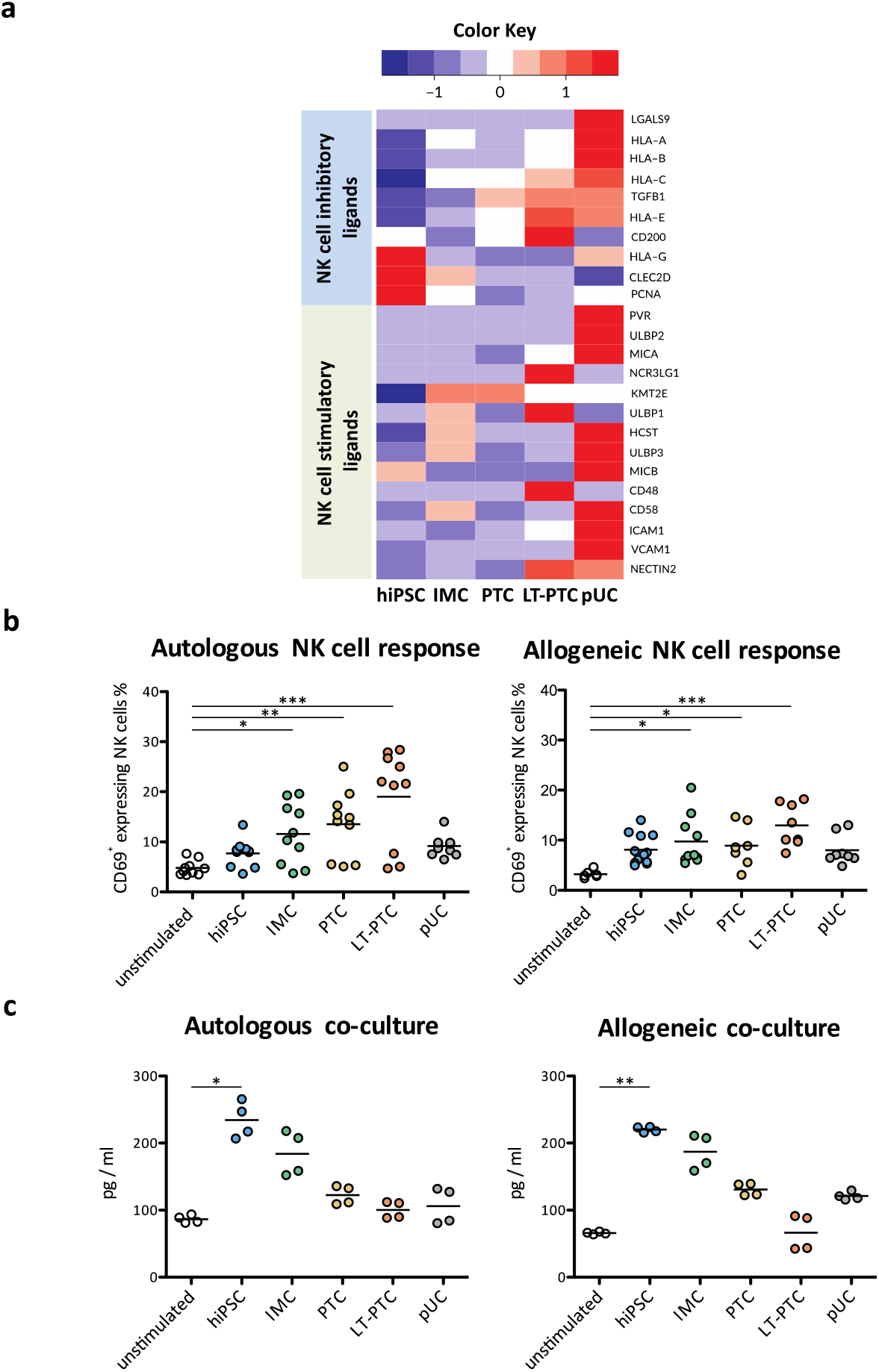
hiPSC and hiPSC-derived renal cells are prone to autologous and allogeneic NK cells. **a)** Transcriptome analysis was used to identify expression intensities of NK cell stimulatory and NK cell inhibitory ligands on hiPSC, IMC, PTC, LT-PTC and pUC for comparison. Three biological IFNγ stimulated replicates were used for each cell type. **b)** Co-cultures were performed with autologous and allogeneic PBMCs from healthy donors. After one day of co-culture, PBMCs were harvested and NK cells were analyzed for CD69 using flow cytometry. NK cell activation was increased when exposed to autologous (n=6-12) and allogeneic (n=10) IMCs, PTCs and LT-PTC. **c)** After 1 day of co-culture the accumulation of IFNγ in the co-culture media was analyzed by multiplex assay. Obtained data revealed significant elevations of IFNγ amounts in co-culture of PBMCs with autologous and allogeneic hiPSC, respectively (n=4). The expression values (FPKM) of each gene (row) are normalized by a row z-score. Statistical analysis was performed using one-way ANOVA followed by Dunn’s post-test. *p < 0,05; **p < 0,01; ***p < 0,001.

To assess the activation status of NK cells, we performed one-way MLR of PBMCs with autologous and allogeneic hiPSC, IMC, PTC, LT-PTC and pUC, respectively. NK cells showed significantly increased levels of activation marker CD69 on the cell surface after 24 hours of exposure to autologous as well as allogeneic IMC, PTC and LT-PTC, respectively (Figure 6b). Tracking of NK cell proliferation of patients with diabetic nephropathy after 7 days of one-way MLR, revealed an undistinguishable response between the experimental groups (Figure S10). In contrast to NK cell activation, analysis of the pro-inflammatory cytokine IFNγ in supernatants revealed the highest levels with autologous as well as allogeneic undifferentiated hiPSC (Figure 6c). The observed NK cell activation and IFNγ response were thus comparable for autologous and allogeneic cell sources. In contrast to previous studies, our analysis revealed only a minor impact of IFNγ pre-stimulation of PSC on the activation status of NK cells (Figure S11).^[35]^

## 3. Discussion

Clinical translation of hiPSC-derived cells requires management of their immunological consequences, locally and systemically, which may depend on the grafted cell type, site and mode of delivery.^[6–8]^ The immunogenicity of hiPSC and hiPSC-derived cells is thus a subject of research and controversy. Autologous hiPSC-derivatives were expected to be tolerated by the host immune system. However, previous studies identified several factors impairing the acceptance of iPSC-derived cells in syngeneic animal hosts, caused by genetic alterations and aberrant gene expression associated with cellular immaturity or cultivation conditions.^[3,24,36,37]^ The genes suspected to provide immune targets include ZG16, HORMAD1, SOX2, OCT4, and GATA3. Although ZG16 expression was detectable in PTC and LT-PTC, GATA3 expression in IMC and PTC, and hiPSC expressed OCT4 and SOX2, neither of these cell types triggered autologous T cell proliferation. Overall, autologous T cells did not respond against hiPSC, IMC, PTC and LT-PTC.

Surprisingly, allogeneic mismatched hiPSC and hiPSC-derived renal cells were also not able to elicit T cell proliferation, even in the presence of preformed allogeneic-specific memory T cells. This was not the case for pUC derived from the same donor as the used hiPSC. In healthy individuals, priming of naïve T cells against foreign HLA-molecules occurs through heterologous immunity and prior exposure to allogeneic antigens, for example due to blood infusion, pregnancy or organ transplantation.^[38]^ Compared to unprimed naïve T cells, memory T cells only require HLA/peptide-TCR engagement without significant costimulatory signals and are less susceptible to conventional immunosuppressive drugs.^[39]^ Thus, preformed allogen-specific memory T cells are the major obstacle in solid organ transplantation (SOT) and may likely play a role for allogeneic hiPSC-based CRT. This role was not supported by our allogeneic one-way MLR data. These data rather indicate that hiPSC-derived cells possess immune-privileged capacities and, in contrast to HLA-identical adult tissue cells, neither reactivated memory T cells, nor did they induce naïve T cell proliferation. We hypothesized that active immune-suppression by hiPSC and hiPSC-derived cells may be responsible for this unexpected result. Indeed, when hiPSC-derived cells were exposed to allogeneic B cell stimulated PBMCs, active immunomodulatory properties of hiPSC, IMC and PTC was observed, but not in the more mature LT-PTC and the terminally differentiated adult pUC.

Since FOXP3^+^ regulatory T cells are essential for immune homeostasis and were shown under certain conditions to play a pivotal role in allogeneic mESC and hESC graft survival, we analyzed the potential of hiPSC and hiPSC-derived renal cells to polarize naïve peripheral CD4^+^ T cells into iTreg.^[40,41]^ Only in the presence of additional T cell stimulators, like SEB or allogeneic B cells (data not shown), co-cultures with hiPSC and IMC did increase the number of T cells with a regulatory phenotype. Thus, T cell mediated acceptance of hiPSC and their renal progenitors was not based on iTreg induction. In conclusion, allogeneic hiPSC, IMC, PTC and LT-PTC exhibit immunogenic potential, but their immune-suppressive capacities, which still remain to be fully elucidated, together with an unattractive immune-phenotype leads to T cell tolerance.

It may be possible that different renal cell identities between pUC and hiPSC-derived renal cells, rather than maturation effects led to the different immune responses. pUC are primary cells, which were cultivated to preferentially expand renal tubular epithelial cells and thus, to be phenotypically close to the hiPSC-derived PTC and LT-PTC.^[42]^ Morphology and transcriptome clustering clearly differentiated the various cell types and confirmed phenotypic stability and functional maturation of LT-PTC. This maturation of hiPSC-derived cells is a necessary requirement to achieve functional equivalency for CRT, however, many hiPSC-derived cells are phenotypically immature.

Our data show that this maturation is associated with changes in the immunogenic potential of the cells. Renal differentiation induced a gradual up-regulation of MHC class I and the antigen-processing machinery. The key mediator of MHC class II, CIITA, is constitutively expressed in antigen-presenting cells (APC) and can be transiently induced by inflammatory stimuli such as IFNγ in non-professional APCs, like primary PTC during renal injury.^[43]^ IFNγ is also a potent inducer of MHC class II in semi-professional APCs, and of MHC class I in somatic cells, affecting the immune-phenotype of cells under pro-inflammatory conditions.^[44,45]^ IFNγ receptor genes IFNGR1 and IFNGR2 are expressed in hiPSC and derived renal cells at similar levels (data not shown), but HLA-DR expression was only inducible in LT-PTC. Overall LT-PTC showed a more mature immune-phenotype compared to PTC. Nevertheless, expression of MHC class I and II molecules in LT-PTC was lower compared to somatic pUC.

Other than T cell responses, NK cell mediated cytotoxicity may play a major role in CRT using hiPSC products.^[4,32,33]^ Here, we revealed that autologous and allogeneic hiPSC-derived renal cells are recognized by NK cells. Autologous as well as allogeneic pUC did not induce NK cell activation, thus, NK susceptibility is not related to allogenicity. Rather it may be caused by reduced expression of MHC class I. Secretion of IFNγ by activated NK cells can stimulate IL-12 production by dendritic cells (DC), which further promote a T helper 1 polarization of naïve T cells due to T-bet induction.^[46]^ Although IFNγ pre-treatment may protect against NK cell-mediated rejection due to upregulation of MHC class I expression, our study showed only a minor impact of IFNγ pre-stimulation of hiPSC on the activation status of NK cells.^[35]^ NK cells trigger an early immune event in organ transplantation, contributing to acute rejection and should be controlled in clinical application of autologous as well as allogeneic hiPSC derived cells.^[47]^ This control requires the analysis of human NK cell responses, preferentially in *in vitro* co-cultivation systems, since common humanized mouse models showed impaired reconstitution with NK- and cytotoxic T cells, which is also seen in allogenized mouse models.^[48,49,50]^ We opted to use fully humanized *in vitro* assays for sensitive detection of immune effects, which could perhaps also be used for individualized preclinical assessment of donor cells and recipient. Furthermore, *in vitro* platforms allow for functionality testing of the therapeutic cells just before their clinical application.

## 4. Conclusion

The therapeutic use of hiPSC-derived renal cells might be a game changer to combat AKI and CKD associated kidney failure and to overcome the high demand for kidney allografts. Our findings show maturation dependent-immunogenicity of hiPSC-derived renal cells, which strongly favors the use of immature tissue specific-cell types due to their strong immunomodulatory capacities even in an inflammatory tissue environment. However, advancing maturation of the hiPSC-derived grafted cells may require control of arising immune-competence and loss of immune-tolerance. This switch must be considered in preclinical assessment platforms of the therapeutic cell product. Furthermore, the immunomodulatory effects of immature hiPSC-derived precursor cells may favorably shape the local graft environment by reducing T cell activation. However, NK cell activation should be taken into consideration. Additionally, a putative risk of this low immunogenicity might be loss of control in case of viral infection or transformation. *In vitro* monitoring and assessment of the dynamic shift between immune-privileges, immunomodulation and immune-maturity stages could thus refine CRT. Finally, the applied example of renal differentiated cells may also be principally relevant for other solid mesoderm-derived tissues, but this will need comparative assessment.

## 5. Experimental Section

### 5.1. Ethics statement

All human cells were obtained following informed consent approved by the Ethics Committee of Charité Universitätsmedizin Berlin, which also covers derivation and use of hiPSC (Approval number EA4/110/10 and 126/2001).

### 5.2. Human cell lines

Two hiPSC-lines BCRTi004-A and BCRTi005-A were used, generated from primary urinary cells (pUC) of two healthy female donors using integration-free Sendai virus (SeV)-technology.^[20,21]^ Conditional immortalized B cell-lines were a gift from Dr. Si-Hong Luu. The generation was performed as described previously.^[51]^

### 5.3. Differentiation of hiPSC into renal cells types

hiPSC were differentiated into IMC and into PTC using a stepwise protocol with slight modifications.^[14]^ Briefly, for IMC-differentiation 4×10^4^ hiPSC / cm^2^ were seeded on Geltrex (Thermo Fisher Scientific)-coated plates in TeSR-E8 media (Stem Cell Technologies) supplemented with Y27632 (Wako Chemicals). After two days, mesendoderm differentiation was induced using 5 μM CHIR9022 (Tocris) dissolved in complete Advanced RPMI (A-RPMI, Thermo Fisher Scientific) supplemented with 1 % GlutaMax (Thermo Fisher Scientific) and 1% Penicillin/Streptomycin solution (Thermo Fisher Scientific). After 35 hours, media was switched to intermediate mesoderm induction media for 72 hours, composed of complete A-RPMI supplemented with 2 μM Retinoic acid (Stemgent) and Fibroblast growth factor 2 (Peprotech). IMCs were cultivated for 7 days in complete A-RPMI to obtain PTC. PTC were cultivated for another 14 days on Geltrex-coated plates to promote their further maturation (LT-PTC). Differentiated cells were harvested for analysis and co-culture assays on day 5 (IMC), day 12 (PTC) and day 26 (LT-PTC) post-induction.

### 5.4. Human primary cells

pUC from healthy donors were isolated as described previously. ^[42]^ After expansion in complete A-RPMI, pUC were cryopreserved in heat inactivated fetal calf serum (FCS, Biochrom) supplemented with 10 % Dimethyl sulfoxide (DMSO, Sigma Aldrich) until use. PBMCs from healthy and diseased donors were isolated using density gradient centrifugation using Bicoll (Merck). After two washing steps in phosphate-buffer saline w/o calcium and magnesium (PBS w/o Ca and Mg, Thermo Fisher Scientific) and optional erythrocyte lysis (Quiagen) was performed and obtained PBMCs were resuspended in co-culture media, consisting of KnockOut-Dulbecco’s modified eagle medium (KO-DMEM, Thermo Fisher Scientific) supplemented with 20 % KnockOut-Serum Replacer (KOSR, Thermo Fisher Scientific) 1% GlutaMax, 0.1 mM non-essential amino acids (NEAA, Thermo Fisher Scientific), 1% β-Mercaptoethanol (Thermo Fisher Scientific) or stored in cryopreservation media at −160 °C until use.

### 5.5. Immunofluorescence staining of adherent cells

Cultured cells were first washed with PBS containing calcium and magnesium. Afterwards, cells were fixed using *Cytofix* (BD Biosciences). Permeabilization was performed using 10 % donkey serum (Milipore), diluted in Perm/Wash buffer (BD). Primary antibody incubation with PAX2 (Life Technologies), LHX1 (Novus), AQP1 (Bio-Techne GmbH) and Na/K-ATPase (Abcam) occurred over night at 4 degrees. After three time washing with Perm/Wash, incubation with fluorescent-labeled secondary antibodies (Thermo Fisher Scientific) was performed in the dark at room temperature. Cells were washed again for three times before the final staining of the nuclei using 4’,6-diamidin-2-phenylindol (DAPI, Sigma-Aldrich). Afterwards, DAPI solution was replaced finally with PBS w Ca and Mg. Cell images were obtained using the Opera Phenix High Content Screening device (Perkin Elmer) and final analysis was performed using Columbus Software (Perkin Elmer).

### 5.6. Transcriptome analysis

Samples were harvested using Gentle Dissociation Reagent (for hiPSC, StemCell Technologies) or Trypsin (Merck) and total RNA was extracted using the Qiagen RNA Mini Kit. cDNA libraries from poly A-tail enriched RNA were prepared from hiPSC, IMC, PTC, LT-PTC and pUC using TruSeq mRNA sample prep kit v.2 (Illumina). Sequence alignment and RNA-Seq analysis: Bcl to fastq conversion was performed using Illumina software (Illumina). Fastq files were aligned against human reference build hg19 provided by the Genome Reference Consortium (GRCh19). Transcript alignment was performed using TopHat. Analysis of differential expression and transcript abundance was performed using Cuffdiff from the Cufflinks analysis package (version 2.1.1). All heatmaps were generated using the gplots package in R-statistical software (version >3.4). The heatmaps picture the mean FPKM for genes (heatmap rows) for different groups (heatmap columns).

### 5.7. Fluorescence labeling of PBMCs

T cell proliferation was tracked using fluorescent dye distribution within daughter cell generation. Up to 10^7^ PBMCs were resuspended in PBS and incubated with either CellTrace Violet (CTV) or CellTrace carboxyfluorescein succinimidyl ester Kit (CFSE, both Thermo Fisher Scientific) in a final concentration of 5 μM.

### 5.8. Flow cytometry

For characterization, hiPSC, IMC, PTC, and LT-PTC were harvested at day 0, 5, 12 and 26 days post-induction, respectively. Collected cells were permeabilized using Perm2 buffer (BD Biosciences) and further blocked in 10 % donkey serum (Merck). Cells were incubated with unlabeled primary antibodies, PAX2, LHX1, AQP1 (Proteintech)) and Na/K-ATPase, respectively, and afterwards stained with secondary conjugated anti-donkey antibodies (Thermo Fisher Scientific). For immune-phenotype analysis of hiPSC, hiPSC-derived renal cells and pUC, conjugated antibodies against HLA-ABC and HLA-DR (BioLegend) were used for cell surface staining. Harvested PBMCs from the supernatants of co-cultures were collected and further stained for vitality (L/D Blue, Invitrogen) and the surface markers CD3, CD4, CD8, CD25, CD40, CD45RA, CD56, CD69, CD80, CD86, CD95, CCR7, HLA-DR (BioLegend, eBiosciences, BD Biosciences, Beckman Coulter Diagnostics). For intracellular staining of FOXP3 and CTLA4 (BD Biosciences), PBMCs were permeabilized using Foxp3 Transcription Buffer Set (eBiosciences). Flow cytometry analysis was performed using a LSR-Fortessa device (BD Biosciences) and results were analyzed with FlowJo 887 (Tree Star). Individual gate settings were performed on the basis of the negative (unstimulated) and positive (SEB stimulated) control.

### 5.9. Immune cell proliferation assay

The stimulator cells hiPSC, IMC, PTC, LT-PTC and pUC were seeded into 24 wells (5×10^4^ per well) in either co-culture media supplemented with Y27632 (hiPSC) or in complete A-RPMI supplemented with Y27632 (renal differentiated cells, pUC). After attachment, cells were stimulated with IFNγ for 48 hours with 25 ng/ml as previously described. ^[17]^ 2×10^5^ CTV labeled PBMCs were added to irradiated (30 Gy) stimulator cells. After 7 days of co-culture, non-adherent PBMCs were harvested and lymphocyte proliferation was assessed by CTV tracking via flow cytometry. For further T cell stimulation, either allogeneic B cells in a 1:10 ratio or SEB (Sigma-Aldrich) in a final concentration of 100 ng/ml was added. For analysis of immunosuppressive capacities, co-cultured hiPSC, IMC, PTC and LT-PTC were used non-irradiated.

### 5.10. Cytokine detection

Supernatants of PBMC co-cultures and mono-cultures of hiPSC, IMC, PTC, LT-PTC and pUC, respectively, were taken on day 1 and 3 and analyzed for TGFβ using ELISA assay (BioLegend) and IFNγ and TNFα using a multiplex-bead based assay (Meso Scale Discovery) according to manufacturer’s protocol. For the determination of total TGF-β, supernatants were treated with acidification solution prior measurement, while the assessment of free active TGFβ was obtained without prior acidification. ELISA samples were measured using the plate reader (SpectraMax 340PC) at 450 nm and at 570 nm to exclude non-specific background staining. Multiplex-samples were detected using the plate reader MESO QuickPlex SQ 120.

### 5.11. Statistical analysis

Results are shown as mean, in certain cases showing mean ± standard error of the mean (SEM). Statistical differences between two groups were assessed using nonparametric Mann-Whitney test. Comparison of more than two groups was performed using Kruskal-Wallis non-parametric one-way analysis of variance (ANOVA) with Dunn’s post-tests. A value of p < 0.05 was considered statistically significant. Graph preparation and statistical analysis was performed using GraphPad Prism 5 (GraphPad Software).

## Supporting information

Supporting information in the supplementary section or from the author.

## Acknowledgement

Thank you to everyone who participated into this work and finalizing the manuscript. We would like to acknowledge the assistance and support by the BIH Stem Cell Core Facility and the Charité / BIH Cytometry Core. We thank additionally Dr. Si-Hong Luu for providing allogeneic B cells from his established B cell bank.

## Author contributions

B.R. and A.K. designed and conceived the experimental setup. B.R. performed the experiments. K.H. supported the experimental design and their execution. N.M. performed analysis of RNAseq data. S.O. provided experimental assistant for cytokine analysis. B.R. and A.K wrote the manuscript. P.R. supervised the project, provided the clinical sample and revised the paper. D.V. mentored the project and revised the paper.

---

**Table 1:**
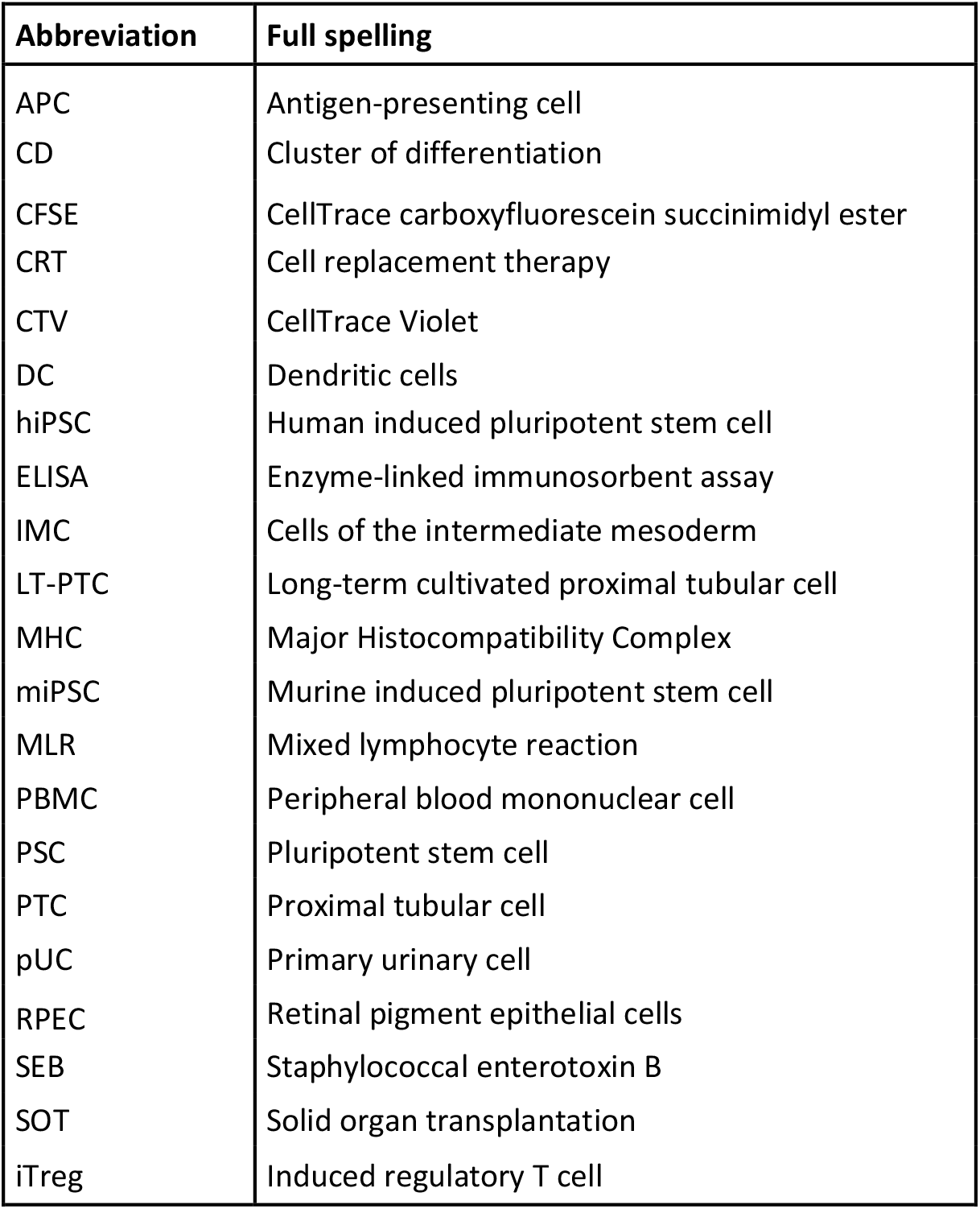
List of abbreviation

| Abbreviation | Full spelling |
| --- | --- |
| APC | Antigen-presenting cell |
| CD | Cluster of differentiation |
| CFSE | CellTrace carboxyfluorescein succinimidyl ester |
| CRT | Cell replacement therapy |
| CTV | CellTrace Violet |
| DC | Dendritic cells |
| hiPSC | Human induced pluripotent stem cell |
| ELISA | Enzyme-linked immunosorbent assay |
| IMC | Cells of the intermediate mesoderm |
| LT-PTC | Long-term cultivated proximal tubular cell |
| MHC | Major Histocompatibility Complex |
| miPSC | Murine induced pluripotent stem cell |
| MLR | Mixed lymphocyte reaction |
| PBMC | Peripheral blood mononuclear cell |
| PSC | Pluripotent stem cell |
| PTC | Proximal tubular cell |
| pUC | Primary urinary cell |
| RPEC | Retinal pigment epithelial cells |
| SEB | Staphylococcal enterotoxin B |
| SOT | Solid organ transplantation |
| iTreg | Induced regulatory T cell |

## Table of contents

hiPSC-derived cells represent a promising approach of functional renal cells suitable for autologous cell-replacement therapies. Yet, the immunogenic propensity is of major concern. Within the described study an immune-suppressive and immune-invisible phenotype of early hiPSC-derived renal cells was detected. In comparison, allogeneic primary cells induced strong T cell reactions. However, hiPSC-derived cells are susceptible to NK cells.

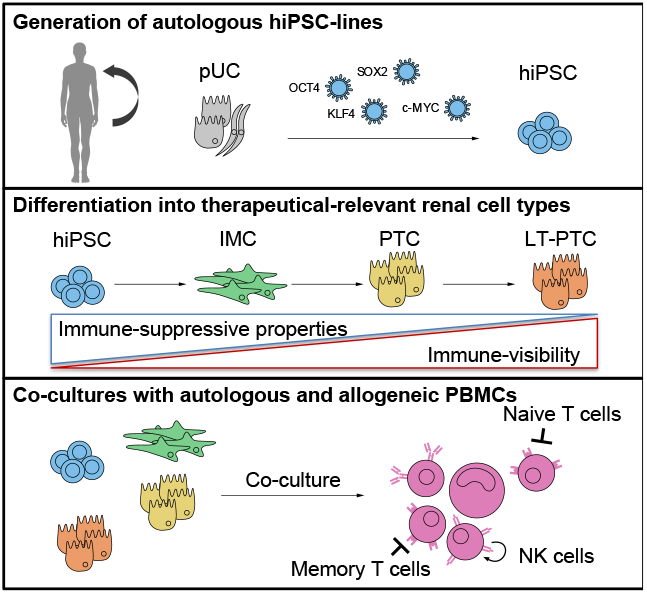

## Supplememtary Information

**Figure S1:**
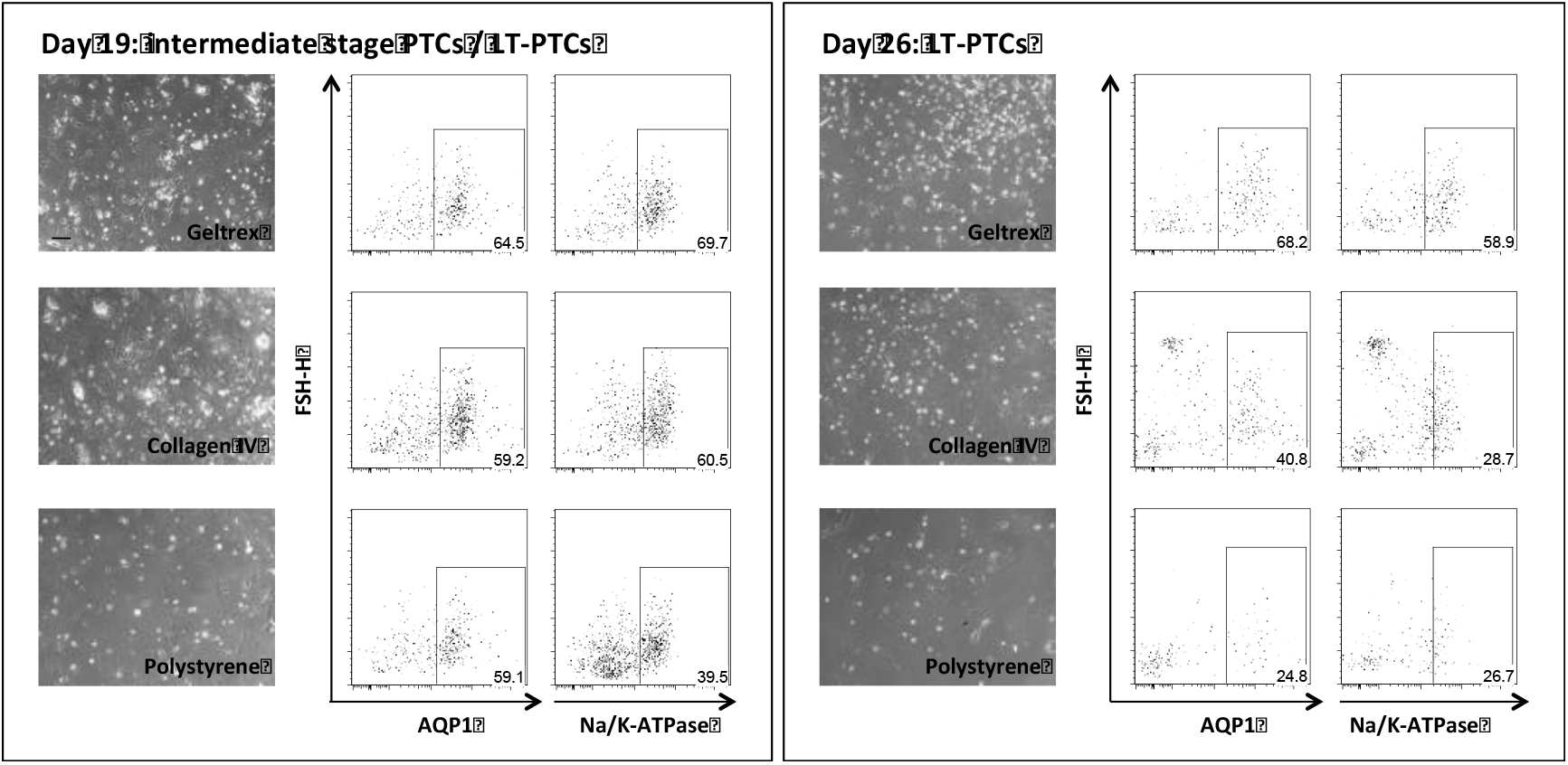
hiPSC-derived PTC can be maintained *in vitro*. After terminal differentiation PTC were cultivated for additional two weeks testing different coating reagents. Using flow cytometry analysis Geltrex was revealed as the optimal matrix for the stable maintenance of PTC to LT-PTC based on AQP1 and Na/K-ATPase expression.

**Figure S2:**
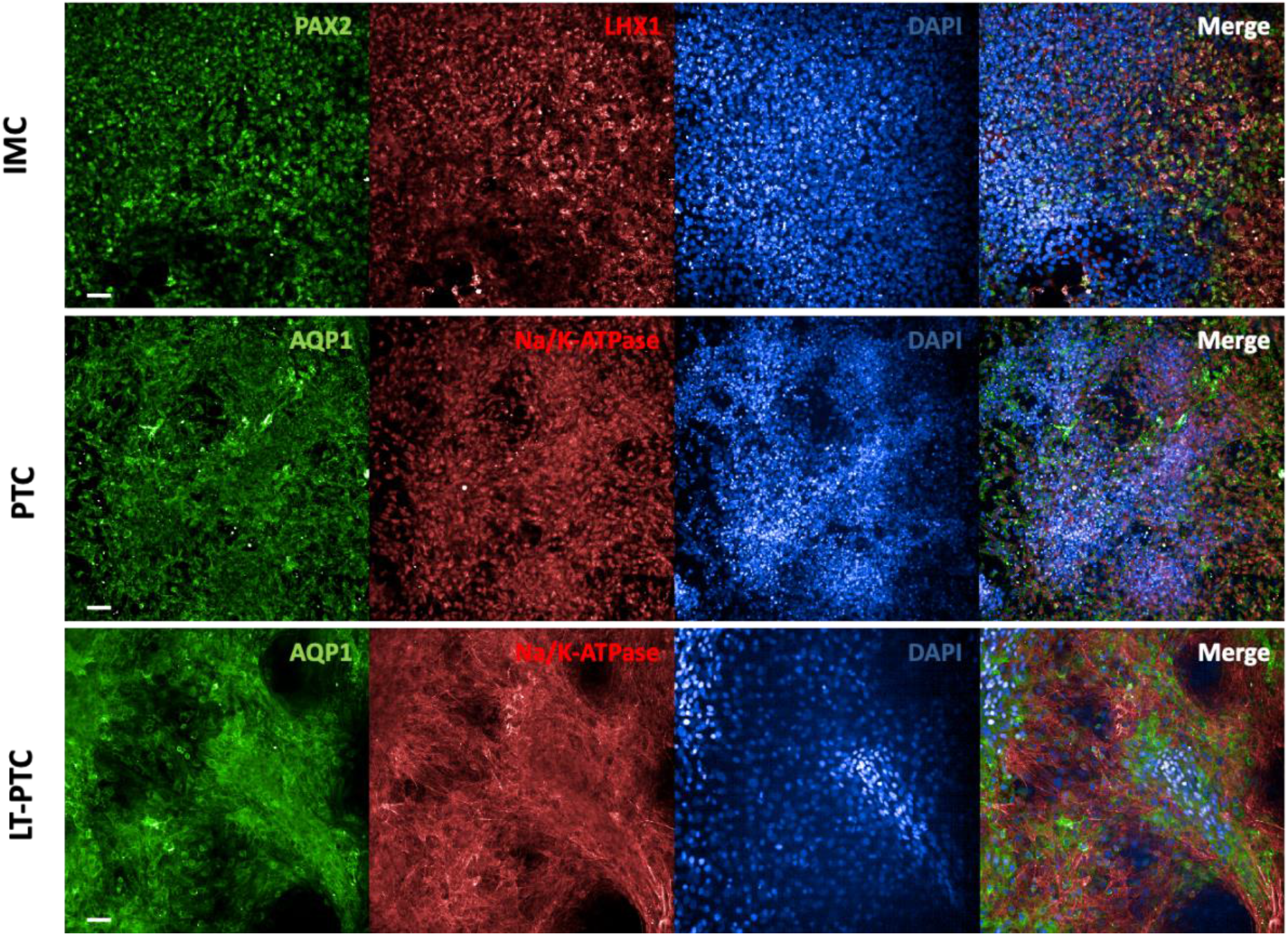
IMC, PTC and LT-PTC show co-localization of stage-specific proteins. Immunofluorescence staining was performed to analyze the localization and co-expression of stage-specific markers in IMC (transcription markers PAX2 and LHX1) and in the proximal tubular cells PTC and LT-PTC (water channel AQP1 and the ion channel Na/K-ATPase). Background staining in the negative control appearing from the secondary antibody was substracted. Scale bar is equivalent to 50 μm.

**Figure S3:**
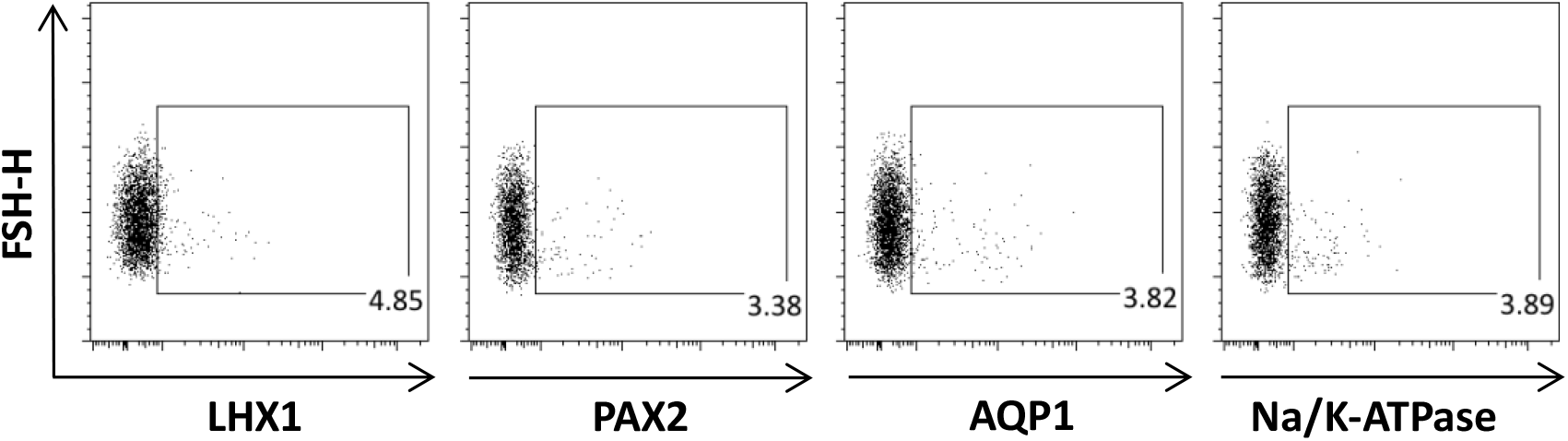
IMC and PTC specific marker are almost not present in hiPSC. Using flow cytometry the expression of IMC and PTC specific marker was tested in hiPSC. Only background expression of the IMC marker PAX2 and LHX1 and PTC marker markers AQP1 and Na/K-ATPase was detectable in hiPSC.

**FigS4:**
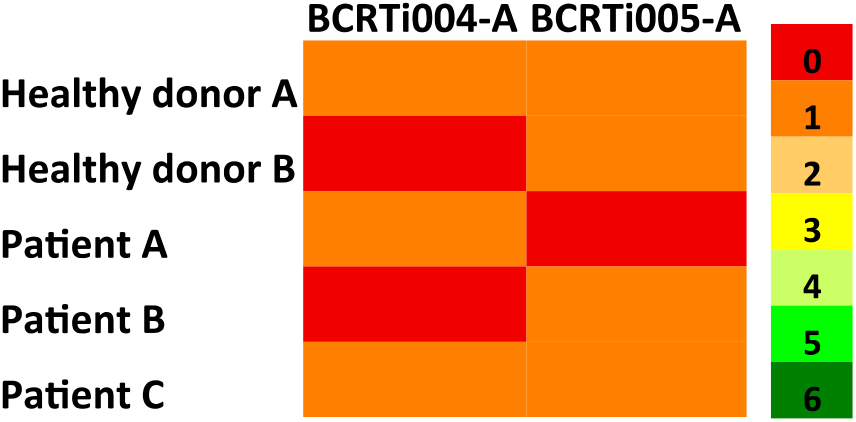
HLA-typing of healthy donors and patients with diabetic nephropathy revealed a maximum match in 1 out of 6 alleles with the hiPSC-lines BCRTi004-A and BCRTi005-A. From two healthy unrelated donors and 3 patients enrolled in the study HLA-types were available. Comparison with the HLA-A-, HLA-B- and HLA-DR-alleles, critical for kidney transplantation, of the hiPSC-lines revealed a maximum overlap of one allele with the allogeneic donors.

**Figure S5:**
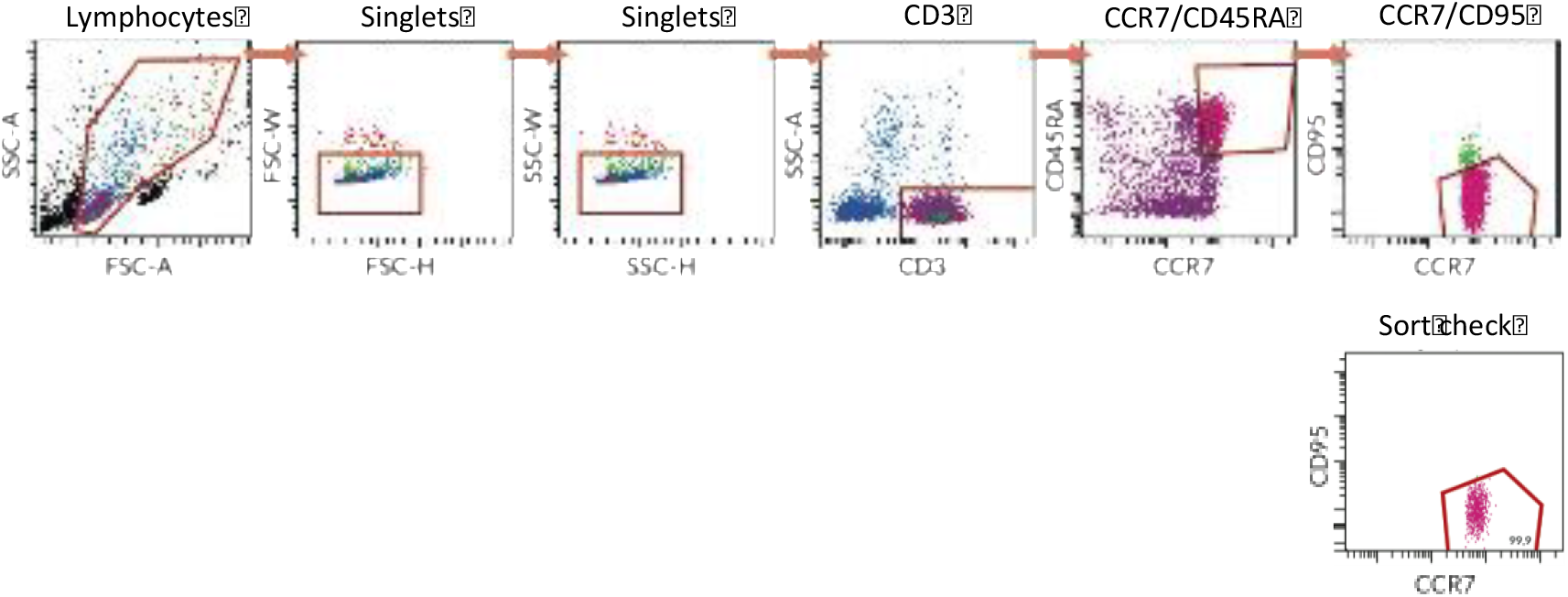
Pure populations of real naïve CD3^+^ T cells were sorted from full PBMCs using FACS. Isolated PBMCs from healthy donors were applied to a FACS device. The subpopulation of single CD3^+^CD45RA^+^CCR7^+^CD95^-^ cells was collected and subsequent sort check revealed a purity of up to 99.9%.

**Fig. S6:**
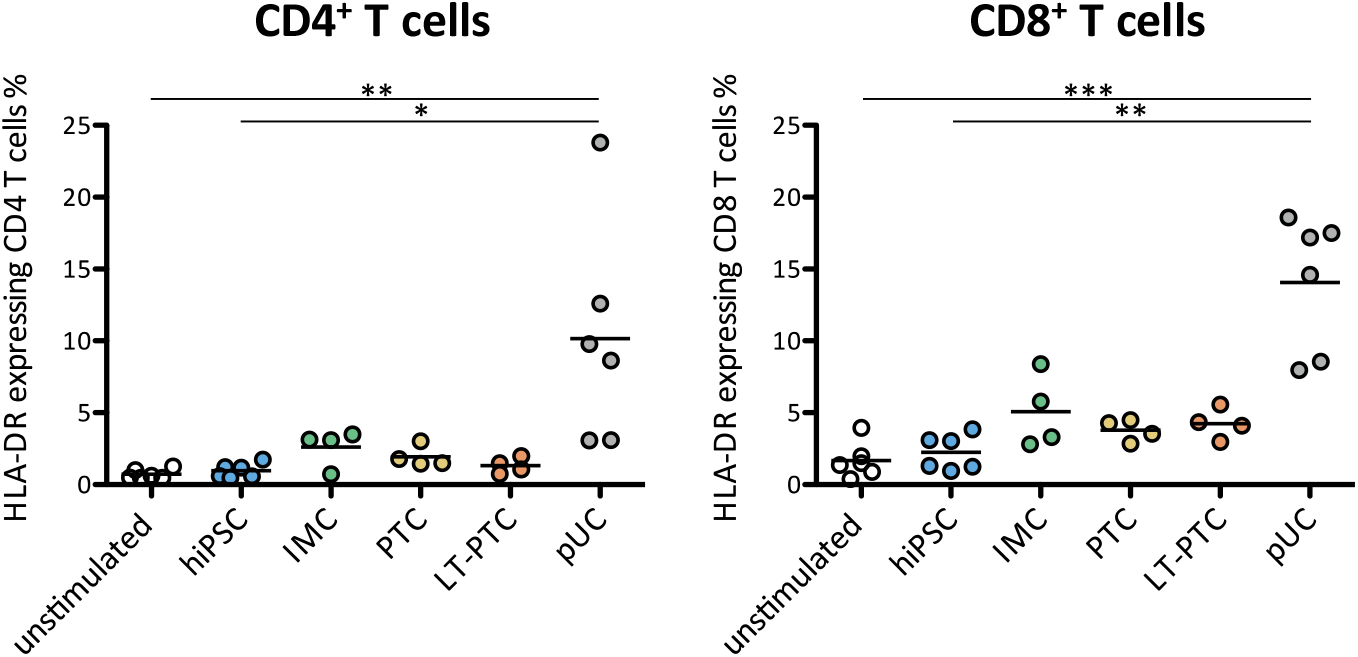
HLA-DR expression is elicited on CD4^+^ and CD8^+^ T cells after co-culture by allogeneic pUC, and not by hiPSC and renal derivatives. Expression of the late activation marker HLA-DR was assessed using flow cytometry on CD4^+^ and CD8^+^ T cells after a co-culture of 7 days with allogeneic hiPSC, renal-derivatives and pUC, respectively. Only allogeneic pUC induced an increase of HLA-DR expression on CD4^+^ and CD8^+^ T cells, whereas hiPSC, and hiPSC-derived renal cells did not trigger activation of allogeneic T cells.

**Fig. S7:**
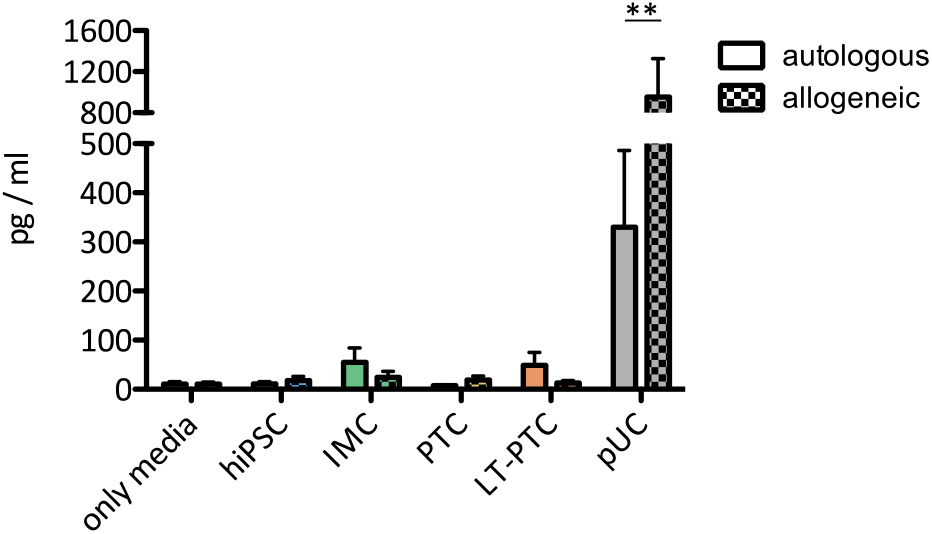
Elevated TNFα release was detected after co-culture of pUC with allogeneic PBMCs. Supernatant on day 3 of all experimental co-culture groups was analyzed for the accumulation of the pro-inflammatory cytokine TNFα via multiplex-assay. In comparison to autologous co-cultures, unstimulated controls and allogeneic co-cultures with hiPSC and renal derivatives, only allogeneic pUC induced increased TNFα release. Statistical analysis was performed using two-way ANOVA followed by Bonferoni’s post-test. **p < 0,01; ***p < 0,001.

**Fig. S8:**
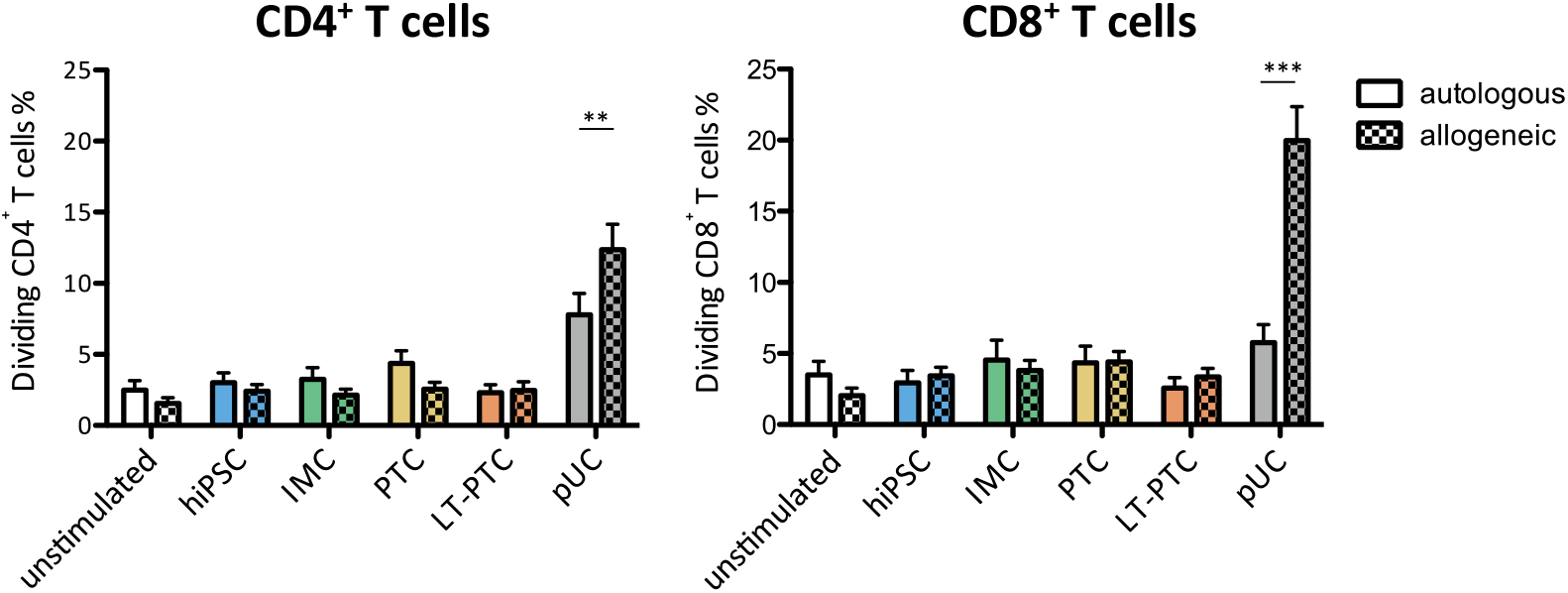
Direct comparison of autologous and allogeneic T cell responses against hiPSC, renal derivatives and pUC revealed only immunogenic capacities of allogeneic pUC, whereas hiPSC-derived stimulators did not elicit T cell proliferation. Autologous and allogeneic CD4^+^ and CD8^+^ T cell proliferation was assessed after 7 days of co-culture with hiPSC, renal derivatives and pUC, respectively. After direct comparison between certain autologous and allogeneic T cell responses, only immunogenicity of allogeneic pUC was observed. In contrast, although sharing the same HLA-type with the pUC, hiPSC and the different renal descendants did not provoke an allogeneic T cell response when comparing to autologous T cell responses as well as to unstimulated control samples. Statistical analysis was performed using two-way ANOVA followed by Bonferoni’s post-test. **p < 0,01; ***p < 0,001.

**Fig. S9:**
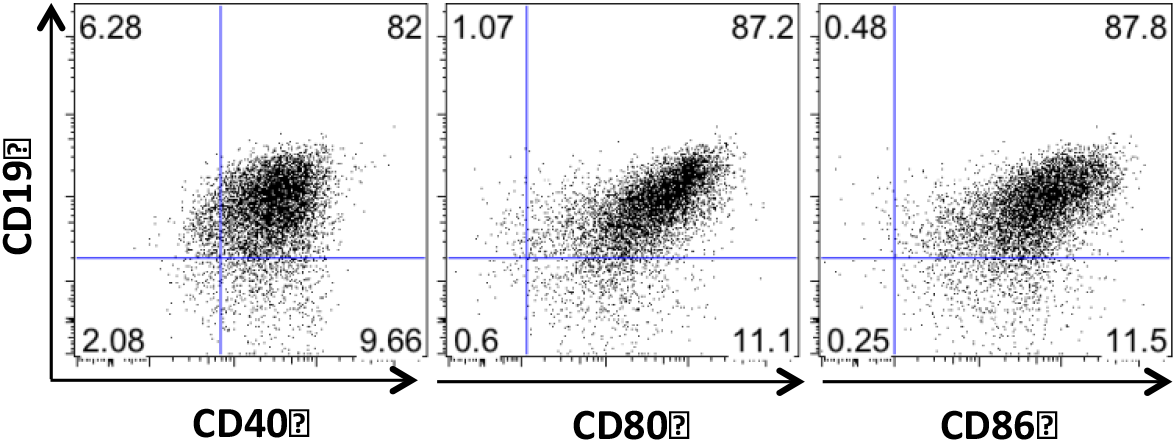
Generated and expanded B cells show an activated immune-phenotype. Freshly thawn B cells were analyzed using flow cytometry. B cells were identified due to expression of CD19. CD19^+^ cells were mainly co-expressing CD40, CD80 and CD86.

**Figure S10:**
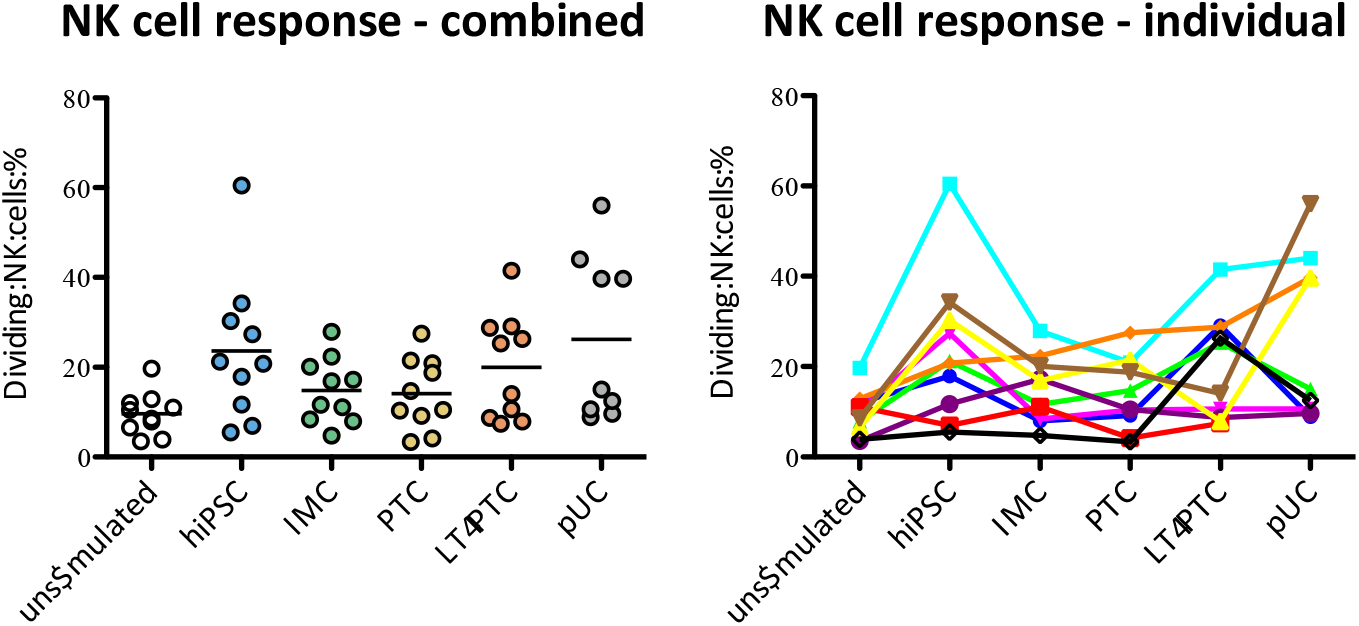
hiPSC, renal derivatives and pUC induced undistinguishable NK cell proliferation in patients with diabetic nephropathy. Proliferation of NK cell was assessed after 7 days of co-culture via flow cytometry. Subsequent statistical analysis using one-way ANOVA testing revealed indistinguishable NK cell response between the different experimental groups.

**Figure S11:**
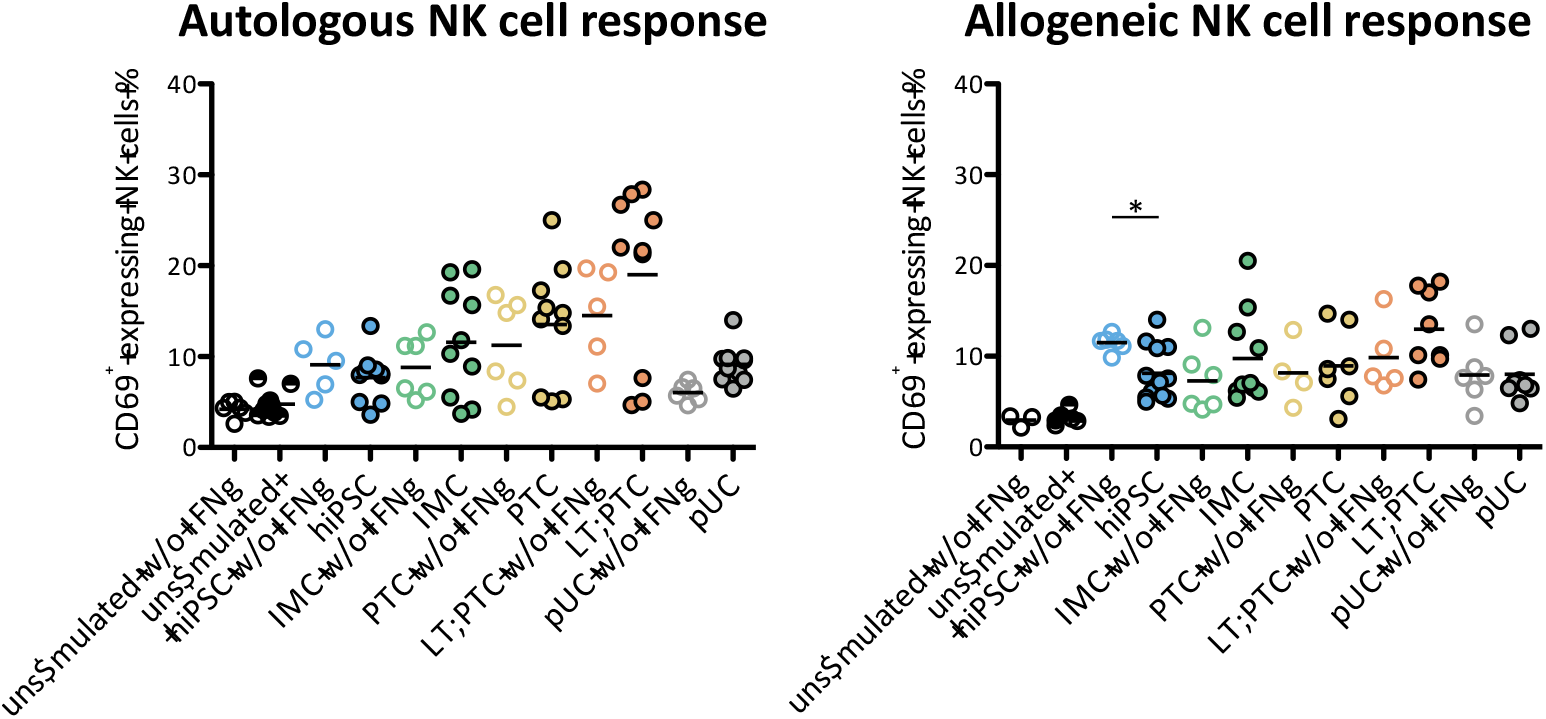
IFNγ pre-stimulated hiPSC elicit less NK cell activation. Using flow cytometry the amount of CD69 expressing NK cells was determined after co-cultivation of PBMCs with unstimulated and IFNγ pre-stimulated autologous and allogeneic hiPSC, IMC, PTC, LT-PTC and pUC, respectively. Comparison of NK cell activation status between unstimulated and IFNγ pre-stimulated samples revealed significant differences in the co-cultures of PBMCs with allogeneic hiPSC. Statistical differences between certain unstimulated and IFNγ pre-stimulated samples were assessed using unpaired Mann-Whitney test. *p < 0,05.

**Table S1:**
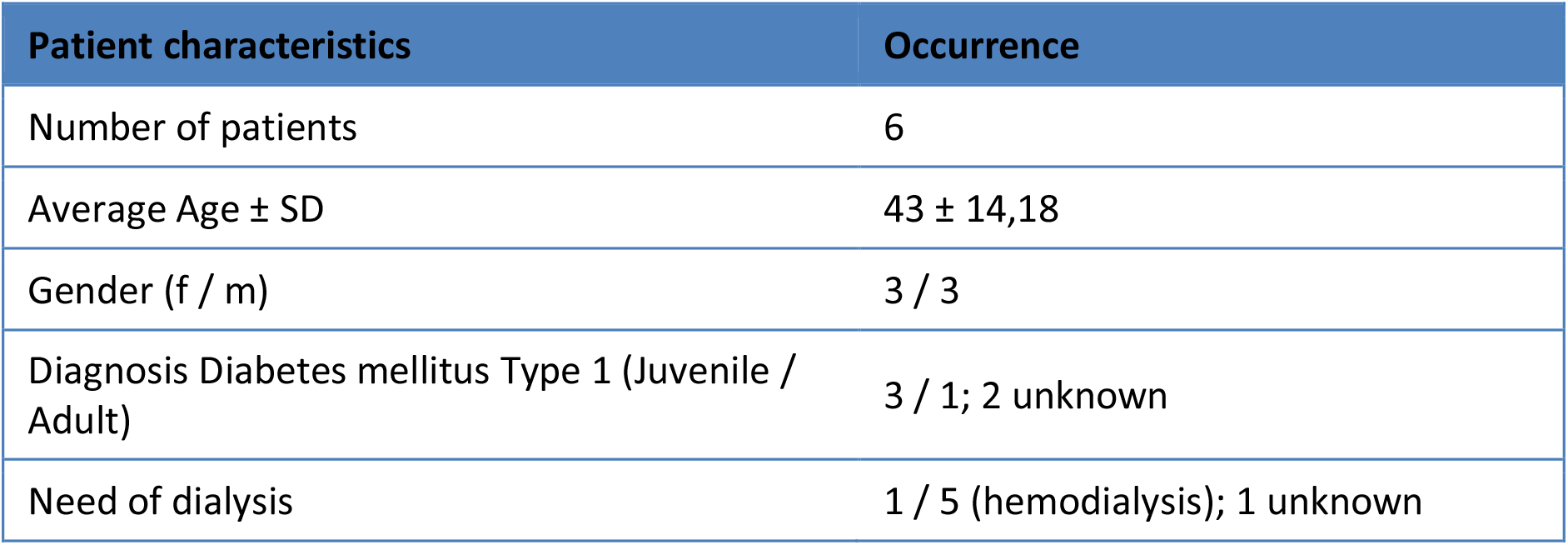
Patient characteristics

