## Supplementary information for "Human iPSC-derived renal cells change their immunogenic properties during maturation: Implications for regenerative therapies"

### Supplemementary Information

Figure S1

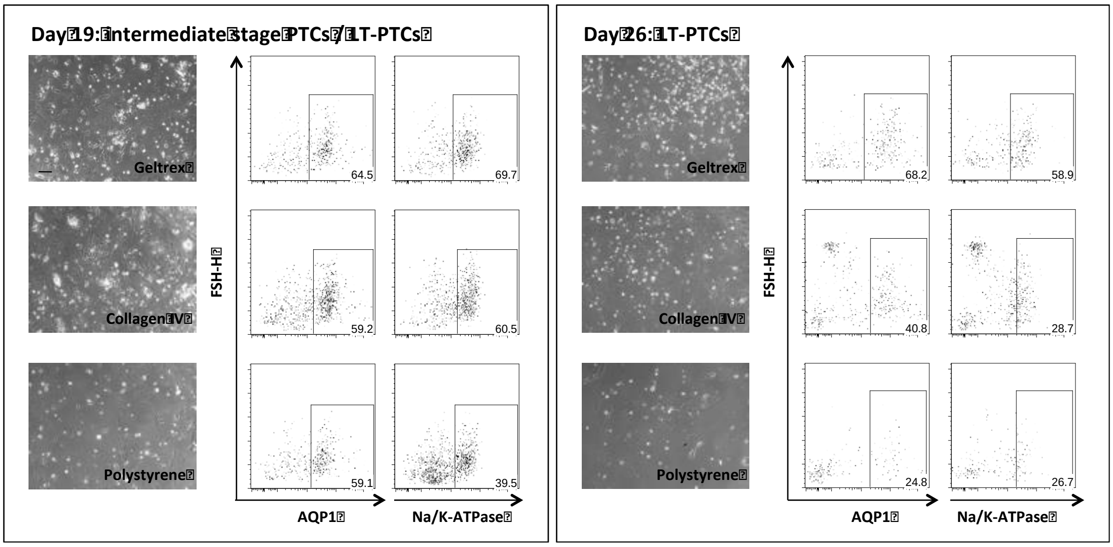


**Figure S1: hiPSC-derived PTC can be maintained *in vitro***. After terminal differentiation PTC were cultivated for additional two weeks testing different coating reagents. Using flow cytometry analysis Geltrex was revealed as the optimal matrix for the stable maintenance of PTC to LT-PTC based on AQP1 and Na/K-ATPase expression.

Figure S2

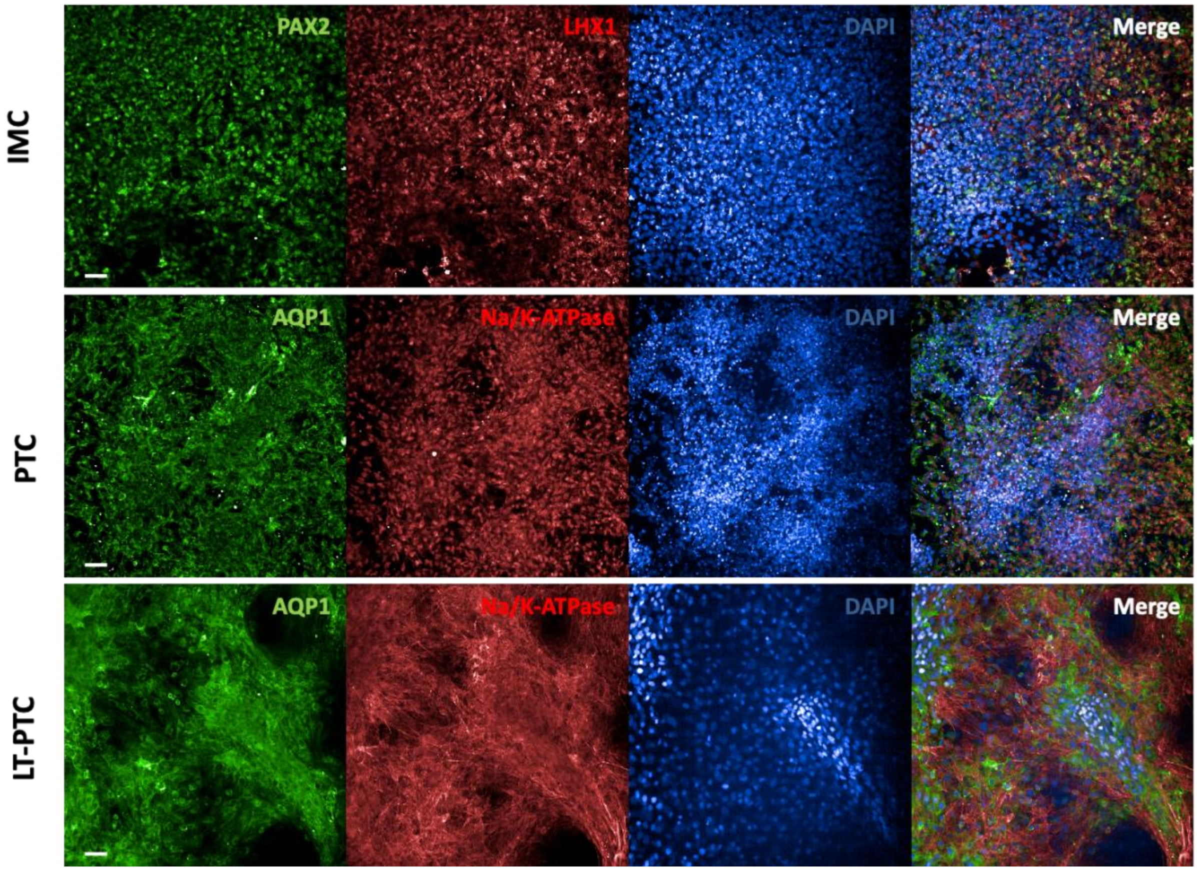


**Figure S2: IMC, PTC and LT-PTC show co-localization of stage-specific proteins.** Immunofluorescence staining was performed to analyze the localization and co-expression of stage-specific markers in IMC (transcription markers PAX2 and LHX1) and in the proximal tubular cells PTC and LT-PTC (water channel AQP1 and the ion channel Na/K-ATPase). Background staining in the negative control appearing from the secondary antibody was substracted. Scale bar is equivalent to 50 µm.

Figure S3

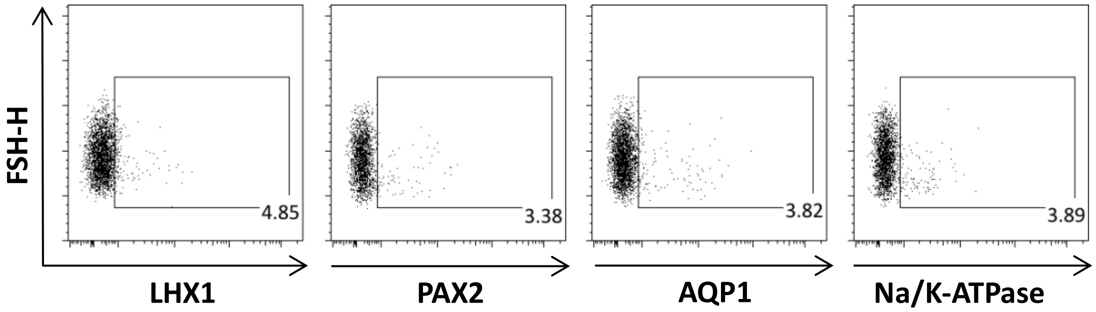


**Figure S3: IMC and PTC specific marker are almost not present in hiPSC.** Using flow cytometry the expression of IMC and PTC specific marker was tested in hiPSC. Only background expression of the IMC marker PAX2 and LHX1 and PTC marker markers AQP1 and Na/K-ATPase was detectable in hiPSC.

#### Figure S4

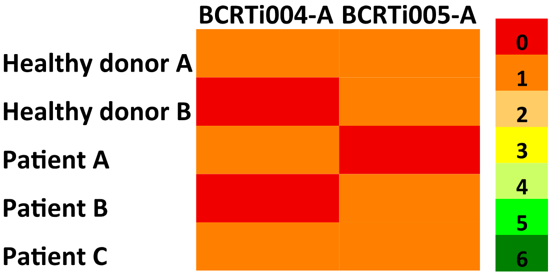


**FigS4: HLA-typing of healthy donors and patients with diabetic nephropathy revealed a maximum match in 1 out of 6 alleles with the hiPSC-lines BCRTi004-A and BCRTi005-A.** From two healthy unrelated donors and 3 patients enrolled in the study HLA-types were available. Comparison with the HLA-A-, HLA-B- and HLA-DR-alleles, critical for kidney transplantation, of the hiPSC-lines revealed a maximum overlap of one allele with the allogeneic donors.

#### Figure S5

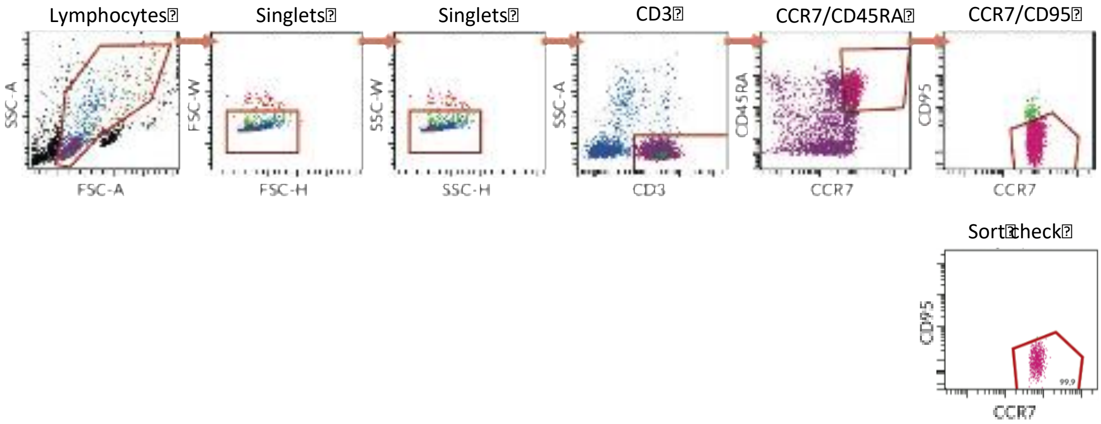


**Figure S5: Pure populations of real naïve $CD3^+$ T cells were sorted from full PBMCs using FACS.** Isolated PBMCs from healthy donors were applied to a FACS device. The subpopulation of single $CD3^+CD45RA^+CCR7^+CD95^-$ cells was collected and subsequent sort check revealed a purity of up to 99.9%.

#### Figure S6

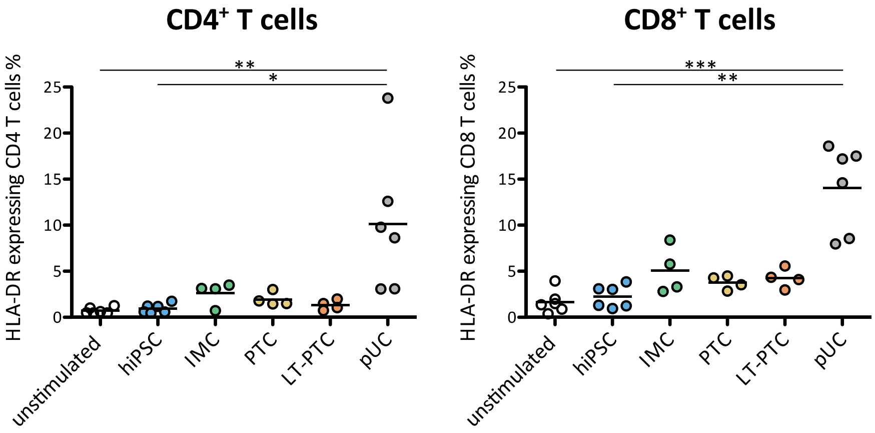

**Fig. S6: HLA-DR expression is elicited on CD4⁺ and CD8⁺ T cells after co-culture by allogeneic pUC, and not by hiPSC and renal derivatives.** Expression of the late activation marker HLA-DR was assessed using flow cytometry on $CD4^+$ and $CD8^+$ T cells after a co-culture of 7 days with allogeneic hiPSC, renal-derivatives and pUC, respectively. Only allogeneic pUC induced an increase of HLA-DR expression on $CD4^+$ and $CD8^+$ T cells, whereas hiPSC, and hiPSC-derived renal cells did not trigger activation of allogeneic T cells.

#### Figure S7

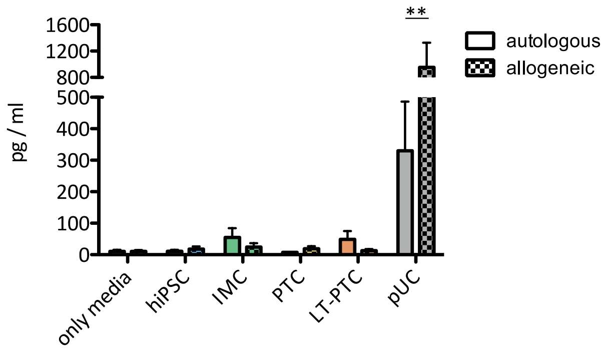


**Fig. S7: Elevated TNF$\alpha$ release was detected after co-culture of pUC with allogeneic PBMCs.** Supernatant on day 3 of all experimental co-culture groups was analyzed for the accumulation of the pro-inflammatory cytokine TNF$\alpha$ via multiplex-assay. In comparison to autologous co-cultures, unstimulated controls and allogeneic co-cultures with hiPSC and renal derivatives, only allogeneic pUC induced increased TNF$\alpha$ release. Statistical analysis was performed using two-way ANOVA followed by Bonferoni´s post-test. **p < 0,01; ***p < 0,001.

#### Figure S8

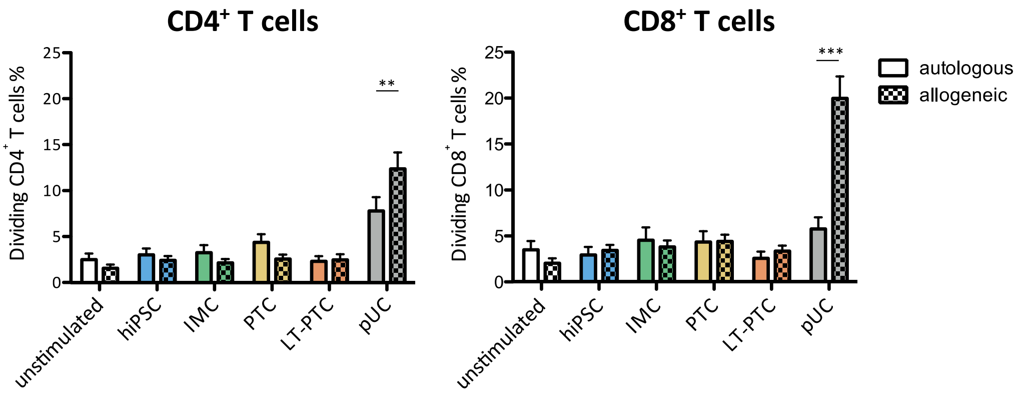


**Fig. S8: Direct comparison of autologous and allogeneic T cell responses against hiPSC, renal derivatives and pUC revealed only immunogenic capacities of allogeneic pUC, whereas hiPSC-derived stimulators did not elicit T cell proliferation.** Autologous and allogeneic $CD4^+$ and $CD8^+$ T cell proliferation was assessed after 7 days of co-culture with hiPSC, renal derivatives and pUC, respectively. After direct comparison between certain autologous and allogeneic T cell responses, only immunogenicity of allogeneic pUC was observed. In contrast, although sharing the same HLA-type with the pUC, hiPSC and the different renal descendants did not provoke an allogeneic T cell response when comparing to autologous T cell responses as well as to unstimulated control samples. Statistical analysis was performed using two-way ANOVA followed by Bonferoni´s post-test. **p < 0,01; ***p < 0,001.

Figure S9

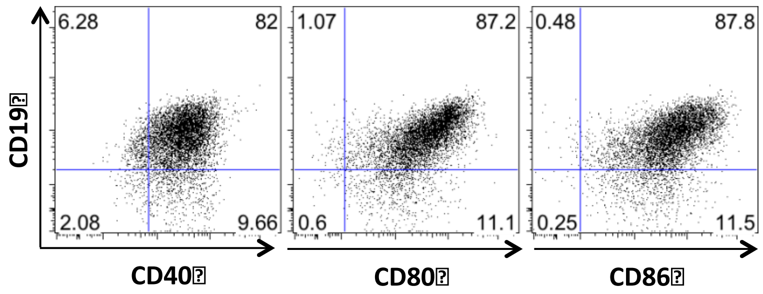


**Fig. S9: Generated and expanded B cells show an activated immune-phenotype.** Freshly thawn B cells were analyzed using flow cytometry. B cells were identified due to expression of CD19. CD19$^{+}$ cells were mainly co-expressing CD40, CD80 and CD86.

Figure S10

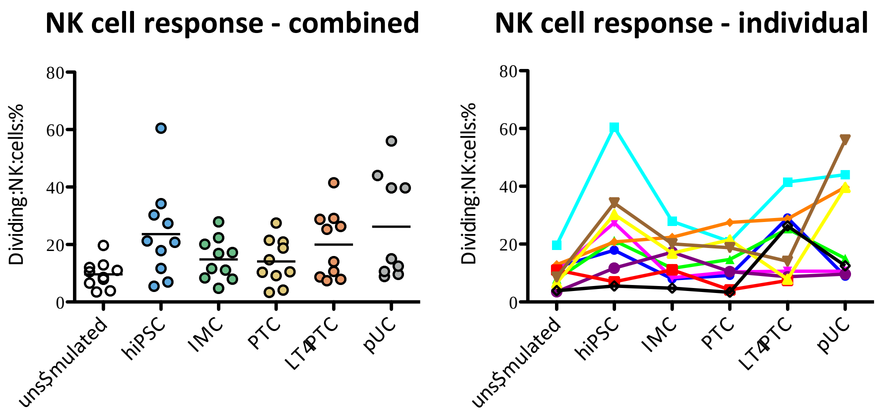


**Fig. S10: hiPSC, renal derivatives and pUC induced undistinguishable NK cell proliferation in patients with diabetic nephropathy.** Proliferation of NK cell was assessed after 7 days of co-culture via flow cytometry. Subsequent statistical analysis using one-way ANOVA testing revealed indistinguishable NK cell response between the different experimental groups.

Figure S11

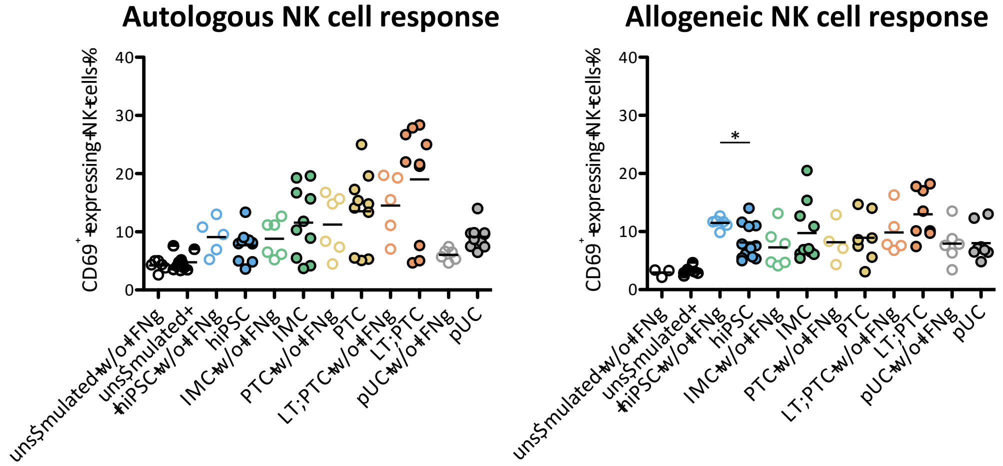

**Figure S11: IFNγ pre-stimulated hiPSC elicit less NK cell activation.** Using flow cytometry the amount of CD69 expressing NK cells was determined after co-cultivation of PBMCs with unstimulated and IFNγ pre-stimulated autologous and allogeneic hiPSC, IMC, PTC, LT-PTC and pUC, respectively. Comparison of NK cell activation status between unstimulated and IFNγ pre-stimulated samples revealed significant differences in the co-cultures of PBMCs with allogeneic hiPSC. Statistical differences between certain unstimulated and IFNγ pre-stimulated samples were assessed using unpaired Mann-Whitney test. *$p < 0{,}05$.

Table S1: Patient characteristics

| **Patient characteristics** | **Occurrence** |
|---|---|
| Number of patients | 6 |
| Average Age ± SD | 43 ± 14,18 |
| Gender (f / m) | 3 / 3 |
| Diagnosis Diabetes mellitus Type 1 (Juvenile / Adult) | 3 / 1; 2 unknown |
| Need of dialysis | 1 / 5 (hemodialysis); 1 unknown |
